# Honey Bees Get Map Coordinates from the Dance

**DOI:** 10.1101/2022.07.27.501756

**Authors:** Zhengwei Wang, Xiuxian Chen, Frank Becker, Uwe Greggers, Stefan Walter, Marleen Werner, Charles R. Gallistel, Randolf Menzel

## Abstract

Honeybees (*Apis mellifera carnica*) communicate the rhumb line to a food source (its direction and distance from the hive) by means of a waggle dance. We ask whether bees recruited by the dance use it only as a flying instruction or also translate it into a location vector in a map-like memory, so that information about spatial relations of environmental cues informs their attempts to find the source. The flights of recruits captured on exiting the hive and released at distant sites were tracked by radar. The recruits performed first a straight flight in the direction of the rhumb line. However, the vector portions of their flights and the ensuing tortuous search portions were strongly and differentially affected by the release site. Searches were biased toward the true location of the food and away from the location specified by translating the rhumb line origin to the release site. We conclude that by following the dance a recruit gets two messages, a polar flying instruction (the rhumb line) and its conversion to Cartesian map coordinates.

## Introduction

Honeybees (*Apis mellifera carnica*) are the only non-human animal that communicate navigational information by a symbolic form of information transfer, the waggle dance, which is performed by successful returning foragers to indicate the direction and distance of the food— in navigational terminology, the rhumb line to food from the hive (1,2). They also use the knowledge about the environment to navigate efficiently and adaptively between multiple locations (3). The learned components consist of a memory of the outbound rhumb line between the hive and previously visited food sources, picture-like memories of the immediate surrounding of the nest and the places where food has previously been found, and memories of the sky-line profiles. Furthermore, they learn the olfactory, gustatory and visual (color, geometric) features of the sources (4-7). Thus, multiple features of the landscape and the properties of previously visited food sources are stored in memory. The question arises whether they are composed in spatial relations and, if so, in what form.

The structure of the forager’s memory of the environment is conceptualized by some authors as multiple unintegrated snapshots. These snapshots are thought to be associated with vectors pointing back to the hive (home vectors) from familiar landmarks or toward locations where food has previously been found (8-10). Others conceive of the memory structure as a cognitive map, from which geometric relations (distance and direction) may be inferred by the navigator (11-13). Here we ask how this memory structure and dance communication systems are linked and what that tells us about the nature of the memory structure.

The danced rhumb line has traditionally been construed as a simple flying instruction. However, a navigator with a map may also infer from it the location of the source (its terminus), in which case the dance may give them access to information on a terrain map—information such as the terrain they should see when flying the vector and location coordinates that would enable them to move toward the indicted location in the search phase following termination of the vector.

We address these questions by applying flight tracking with harmonic radar. This technique has been successfully used to prove symbolic information transfer in the waggle dance (14). The recruits started not only at the hive entrance, but in a catch-and-release design also at other sites within the explored area around the hive.

## Results

### Experimental design and flight trajectories

Recruited bees were observed while they attended dancers that advertised for an artificial feeder at a location within the hive’s foraging territory (denoted by F, see Fig. 1). When the recruit appeared at the hive entrance a radar transponder was glued to its thorax. After release (either at the hive or at a distant release site) her flight trajectory was followed by harmonic radar with radar fixes every 3 s. Recruits had never found food at F (including natural food, see Methods). Therefore, they did not have vectors pointing toward F from surrounding landmarks nor did they have associated horizon profiles that would attract them in its direction from elsewhere in their foraging territory.

**Fig. 1.**
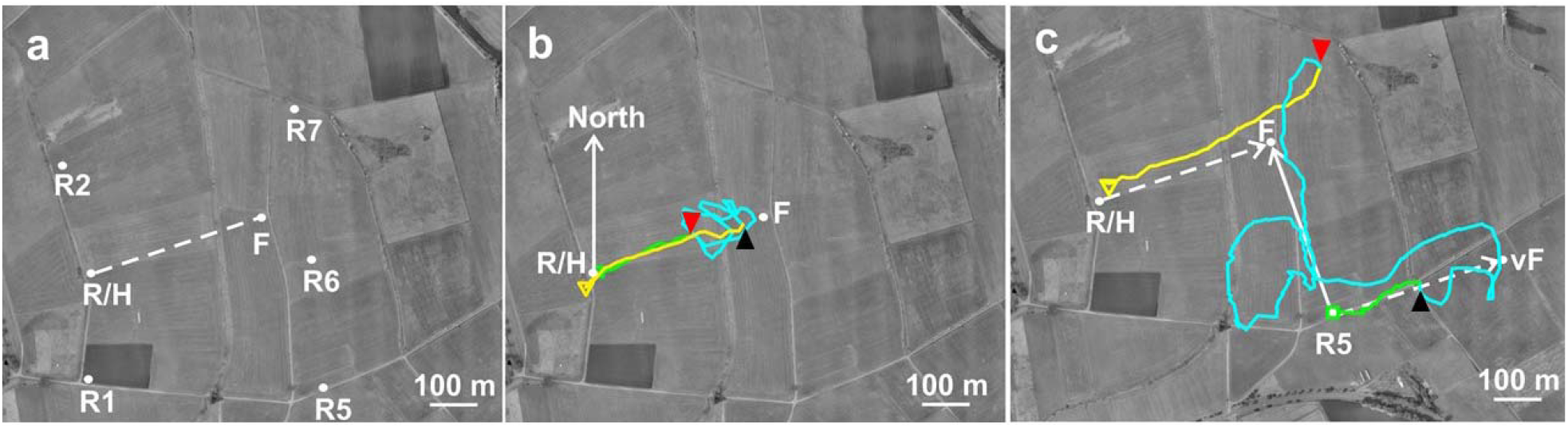
Experimental design and flight trajectories. **a**: Experimental area. R/H: radar and hive location; F: the feeder for the dancing bees; release sites: R/H (hive release), R1, R2, R5, R6, R7. The dotted line gives the polar vector (rhumb line) to F communicated by the dancer (range: 388 m, compass bearing: 71°). **b**: Flight trajectory of a recruit released at the hive. The green square marks the beginning of the flight and the yellow triangle the end of the flight. The black up-pointing arrow head indicates the transition from the outbound vector flight to the search flight, and the red down-pointing arrow head the transition from the search flight to the straight homing flight. **c**: Flight trajectory of a recruit released at R5. Notice the initial vector flight towards the virtual feeder location vF (green), the search flight (blue) composed of the return flight to the release site and the flight composed of a circle and a flight toward the real feeder F, and the homing flight (yellow). The dashed arrows mark the rhumb lines toward F from R/H and from R5 to vF. The dolid arrow from R5 to F marks the shortcut to the real feeder.

Both hive-released recruits and displaced recruits first performed a straight flight after departing from the release site (vector flight), then a tortuous search prior to returning to the hive (Fig. 1 b, c, SF1 a, b). The transition from the vector to the search flight and from the search flight to the homing flight was identified by a sudden turn between two consecutive radar fixes of > 60° after a long straight flight from the release site or immediately preceding a straight return flight (see Methods). Recruits were released not only at the hive entrance but also at five other locations (Fig. 1 a, c; R1, R2, R5, R6, R7). Homing flights were not considered further here because all recruits returned successfully home along direct flights at the end of their search. By design, the danced location had no features indicative of food; for which reason, most recruits released at the hive failed to find it.

### Release-Site Dependent Perturbations of the Vector Flights

We parameterized the outbound vector by i) the compass bearing of the line from the first (f_1_) to the terminal fix (f_t_) in the vector fix sequence, ii) the length of this line, iii) the speed with which the outbound vector was completed [length/(*t*_f_t_–*t*_f_1_)], and iv) its straightness [length/(sum over inter-fix segment lengths)]. Releases from sites other than the hive substantially altered these parameters and the locations relative to the vF where the outbound vector’s terminated. The patterns of alteration differed dramatically depending on the release site, hence, on the terrain the recruit observed during the vector flight (Figs. 2 and 3). Therefore, these effects cannot be attributed to any factor that would be the same for any displaced release site, such as a failure to observe an expected horizon profile on setting out on the vector. Control recruits from another far distant hive released at R2 and R5 mostly flew around in the immediate vicinity of the release site and then disappeared (Fig. 4, bottom two panels). Their “outbound” vector flights were few and short.

**Fig. 2.**
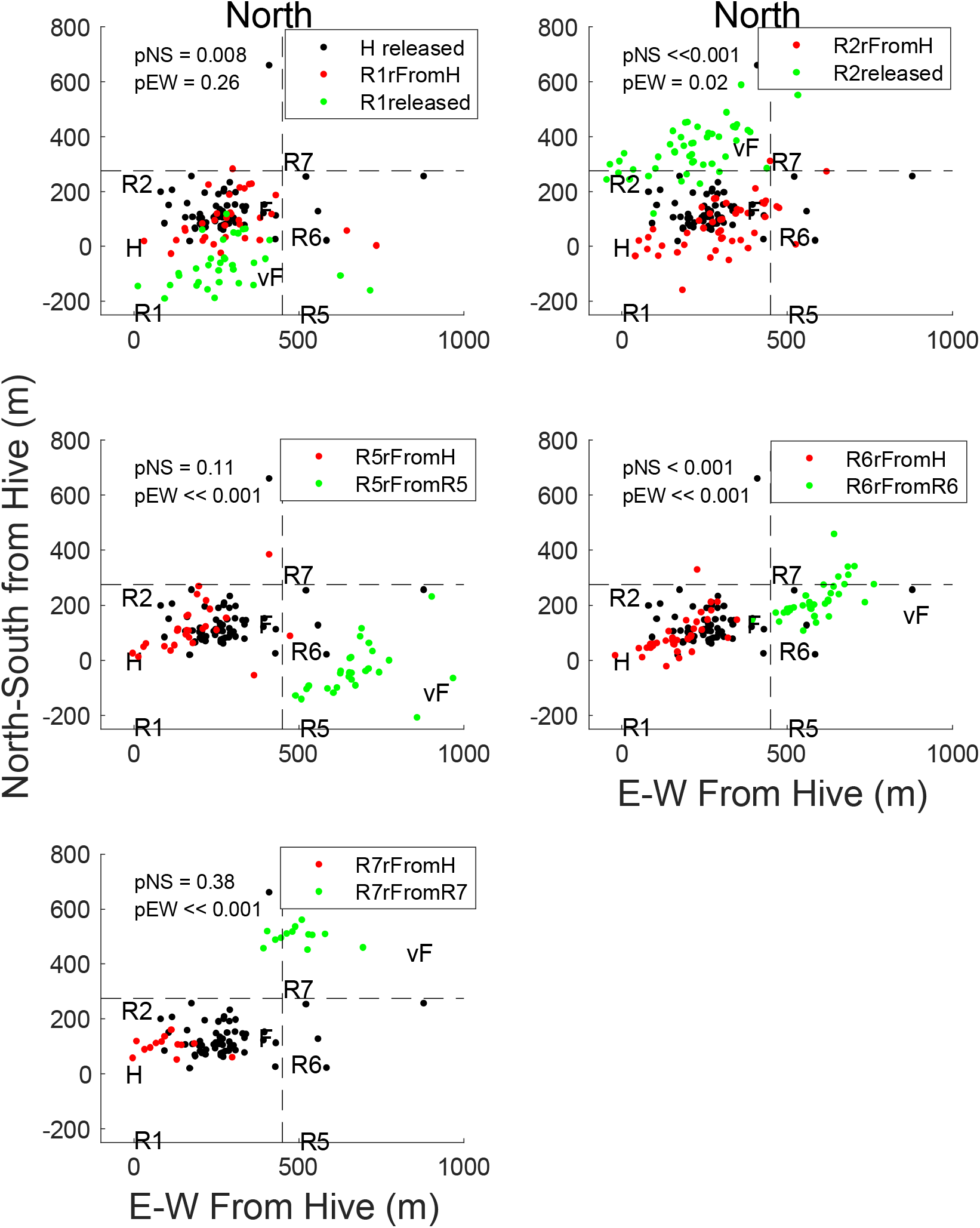
Endpoints of the vector flights from the different release sites shown in relation to the feeder F and the virtual feeder vF. The black dots (repeated in each panel) give the vector endpoints of recruits released at the hive (H), green dots the vector endpoints of the recruits released at the particular release site, and red dots the green dots shifted such that the release site is located at the hive. The shift enables visual comparison of the observed distribution (red dots) to the distribution expected on the rhumb-line hypothesis (black dots). The p values are from Kolmogorov-Smirnov 2-sample tests comparing the latitudinal distribution (north-south coordinates) of the red dots to the latitudinal distribution of the black dots, and likewise for the longitudinal distributions. Note that the pattern of the differences in vector terminations depends on the release site.

**Fig 3.**
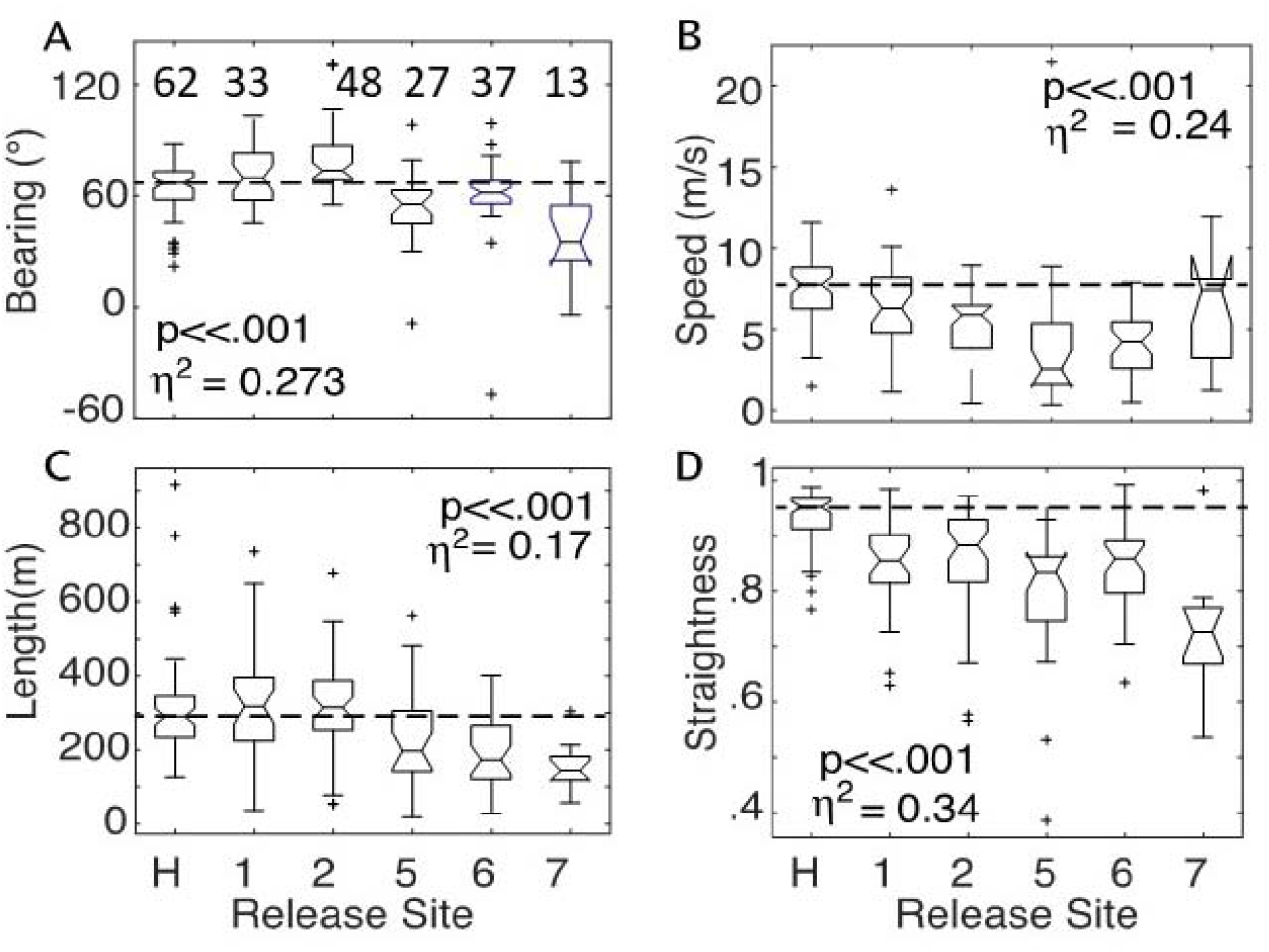
**Effects of release site on parameters of the outbound vector flights** (box plots from one-way ANOVAs). Dashed horizontal lines are at medians for hive-released bees. p = p returned by one-way ANOVA. η ^2^ = effect size (fraction of the variance accounted for by the variation in release site). Numbers above the boxes are the n’s. A. As expected from prior work (11) displaced recruits flew in roughly the same direction as the hive released bees (dashed line), but with systematic release-site dependent deviations. B. Regardless of release site, displaced recruits flew more slowly. C. Recruits released east of F aborted (shortened) their vector flights. D. Regardless of release site, displaced recruits flew more crooked vectors (p<<.001, see comparison tables ST1-4 and SF1 a, b in the Supplementary Material).

**Fig. 4.**
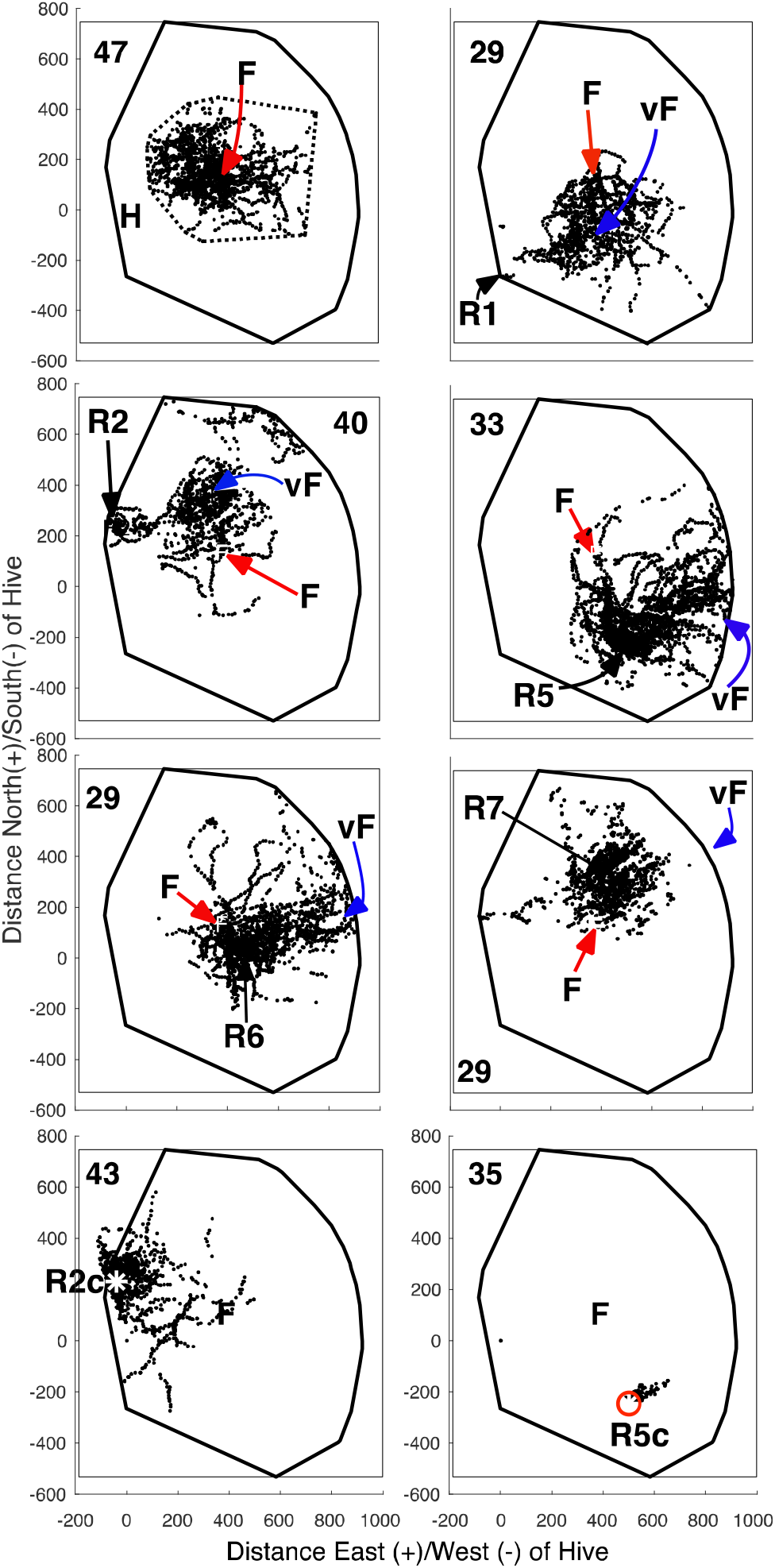
Search-phase fixes for 8 release groups plotted inside the convex hull of the fixes from the 6 groups captured at H. Release site F and vF indicated with bold letters. The bottom two panels (R2c & R5c) are the fixes from two control groups captured on exiting a distant hive; hence, unfamiliar with the territory. The number in a corner of a panel is the number of recruits in the group. The number of fixes contributed by a recruit varied from 4 to 369. The hive-release panel (top left) shows the convex hull of the hive-released searches (dotted outline). It is centered on F. Note that many of the fix patterns for displaced groups are not centered on the vF (e.g., R5, R6, R7).

The site-dependent disruptions of the outbound vector flights and their dependence on familiarity with the test area implies that a recruit’s construal of the dance gives it access to information on a terrain map about what it should and should not see while flying the rhumb line.

### Search Phase Statistics

There is an extensive experimental and theoretical literature indicating that searches efficiently survey terrain centered on an initial estimate of the location of a sought-for goal (e.g., the nest, for a homing ant) (15-17). On the rhumb-line-only hypothesis about a recruit’s interpretation of the dance, a displaced recruit’s estimate of where it should center its search is the terminus of the H->F vector when the origin of this polar vector has been translated to the release site. We call these termini the virtual food locations (vF).

On the rhumb-line only hypothesis, the distributions of fixes from the displaced recruit groups should look like the distribution from the hive-released recruits but centered on the vFs rather than on F. Fig. 4 plots the search-phase fixes from the bees in 8 groups (6 released in their foraging territory and the 2 control groups composed of recruits unfamiliar with the foraging territory—see Methods). It is obvious that the rhumb-line only hypothesis fails at the group level; the distributions of search fixes from displaced recruits do not look like the translated search fixes of hive-released recruits.

As may be seen in the publicly available movies of the individual search-fix sequences (<u>OSF</u>), the rhumb-line-only hypothesis also fails at the within-group level and the within-search level. *Within-group failure:* Different recruits released at the same site make very different searches. *Within-search failure:* In different phases of its search, a recruit often directs it to different locations (F, vF and the release site, RS, see movies). Thus, the assumption that a search has a single center is often false; the early phase of a search may be directed to vF, a later phase back to RS and a still later phase (or phases) to F (see for example, Figure 1c)

On the cognitive map hypothesis, a recruit infers approximate map coordinates for F from the danced rhumb-line by polar-to-Cartesian conversion. If it can also determine approximately where it is during its search, then it should direct some part of its search toward F. How accurately it does so will depend on the accuracy of coordinates inferred for F and on the accuracy of its estimate of its current location. Inaccurately set courses will fail to bring it close to the intended goal.

The simplest test of the rhumb-line-only hypothesis vs the hypothesis that recruits also infer the map coordinates of F is to compare the closest approache (denoted CA) to F and vF. In Fig. 5a, we compare them by computing CA_F_–CA_vF_ for each search. The sign of this difference tells us which it got closer to, F (+) or vF (–). The absolute value, |CA_F_–CA_vF_|, tells us how much closer. Substantial fractions of the recruits in groups R1, R2, R5, R6 and R7 got closer to F than to vF (Figure 5a)—17%, 28% 45%, 69% and 94%, respectively. In the groups released at R6 & R7, 24% and 38% of the bees got more than 200 meters closer. Moreover, a majority of the closest approaches to F fell within the range of closest approaches made by recruits released at the hive (Fig. 5b).

**Fig. 5.**
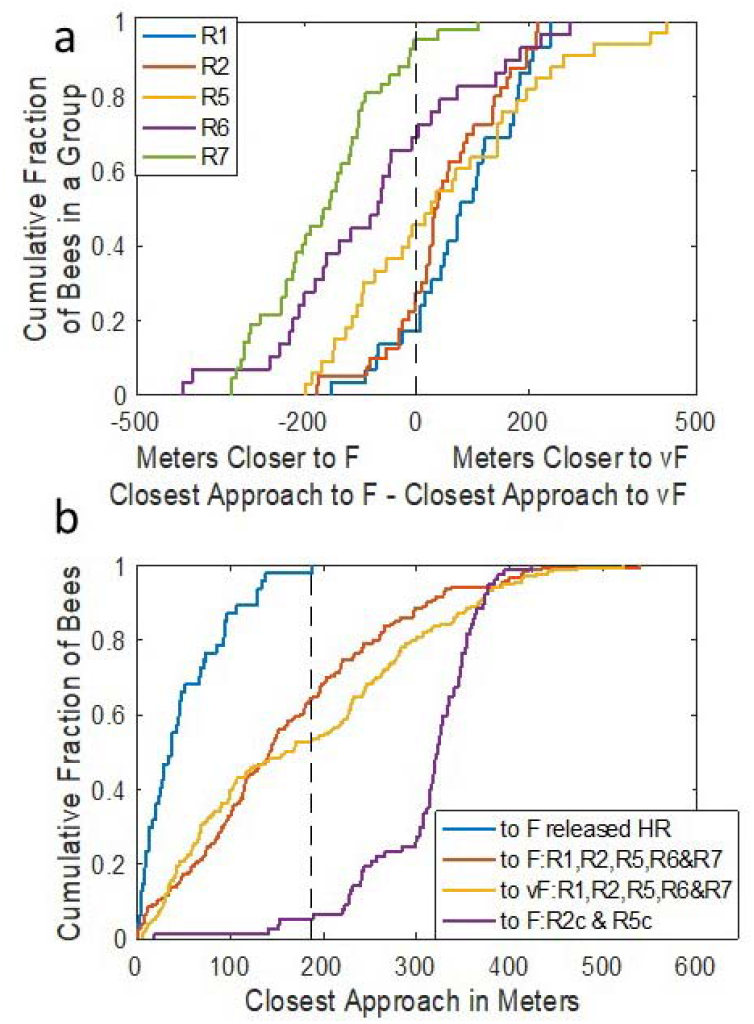
a Cumulative distributions (CDFs) of the bee-by-bee difference between a recruit’s closest approach to F and its closest approach to vF; one CDF for each displaced release group. Each step in a CDF is from a different bee. The fraction on the y axis at the top of a step is the fraction of the differences less than or equal to the value on the x axis. The portion of a CDF to the left of the vertical dashed line at 0 is the proportion of bees in a release group that came closer to F than to vF. (For the n’s, see Fig 7 or count the steps in the plot). b: The CDFs of the closest approaches to F in hive-released recruits (HR, blue), to F in displaced recruits released in familiar territory (red), to vF by displaced recruits released in familiar territory (yellow) and by control recruits from a far distant hive released at the same sites (R2 & R5) as two of the groups from the local hive (purple). The proportion of the yellow, red and purple plots to the left of the vertical dashed line are the proportions of recruits whose closest approach to F was within the range of the hive-released recruits’ closest approaches.

However, F was not the only goal toward which portions of a search were directed. Most displaced recruits flew toward vF initially (Fig. SF1a) and approached it at one point or another in their search, sometimes more than once (see, for example, Fig 7b). Many recruits released to the east of F, hence far to the east of the hive (at R5, R6 and R7, see Fig. 1 a), also went back to their release site at one point or another during their search (Fig 1c), sometimes more than once (Fig. 7b). By contrast, recruits released at H, R1 and R2 centered their searches far to the east of their release site, at roughly the longitude of their vF. These recruits returned to or never left the release site much more rarely (see SF2).

**Fig. 6.**
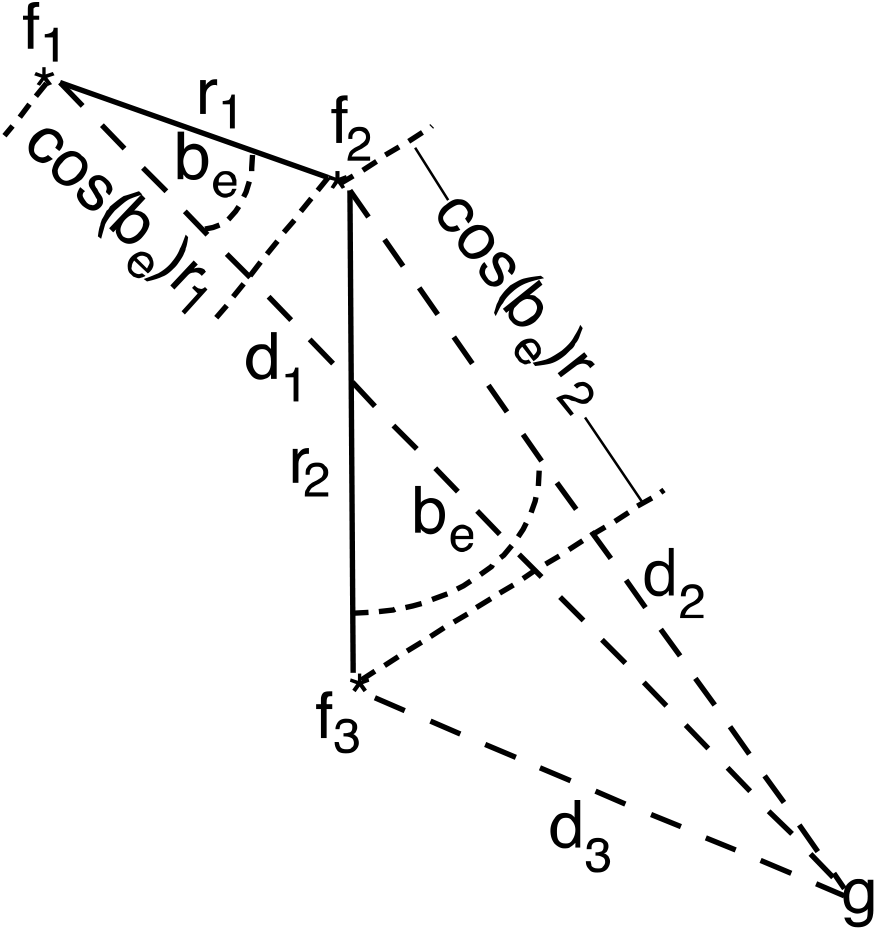
The quantities in the computation of G, the tendency to gravitate toward a target location. g: b_e_ is the bearing error of a fix-fix segment (the angular distance of the bearing of the line between the two fixes from the bearing of g from the first of the two fixes); r_i_ is the fix-fix distance; d_i_ is the distance to g. The progress toward g during a given segment i is cos(b_e_)r_i_.

**Fig. 7.**
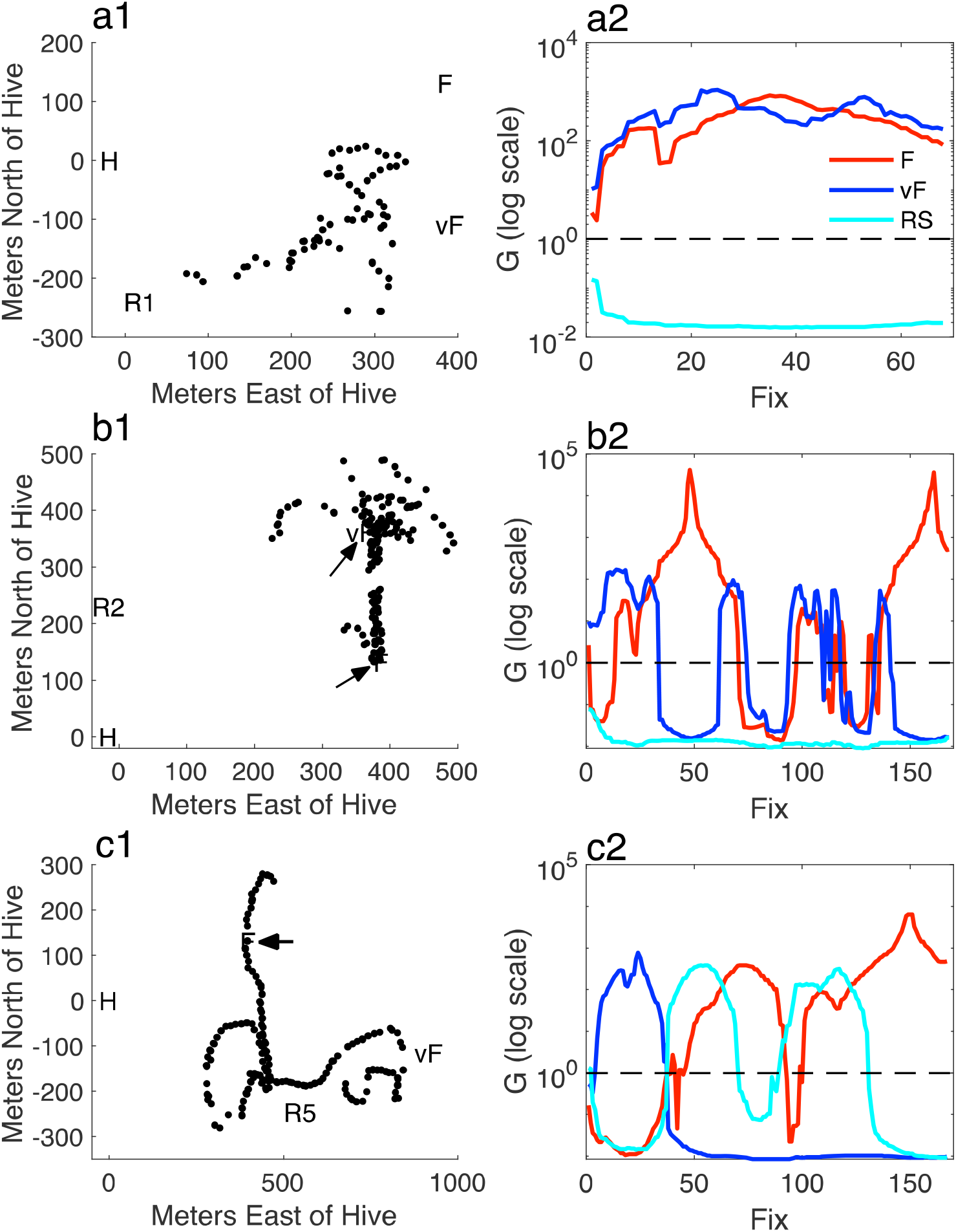
Three illustrative search-fix sequences (panels on left) and the fix-by-fix plots of G_F_ (red), G_vF_ (blue) and G_RS_ (cyan) in the panels on the right. These gravitations are plotted as a function of the successive fixes in the search sequence a: Released at R1. b: Released at R2. Arrows indicate F and vF, which are mostly obscured by fix marks. The recruit flew twice directly from vF to F. c: Released at R5. Arrow indicates F. The recruit went first to vF, then back to NW of R5, then due north toward F, then back to NW of R5, then due north directly over F (see Figure 1c for this same trajectory).

On the map hypothesis, the terrain seen by recruits flying from H, R1 and R2 confirmed that they were flying in the general direction of F. By contrast, the terrain seen by recruits flying east toward vF from R5, R6 and R7 implied that they were flying away from F. This difference in the map-based implications of the terrain traversed by their vector flights explains why the former groups completed their vector flights and then made searches that straddled the longitude of their vFs, while the latter groups aborted their vector flights and searched back toward the west—with the result that the distributions of their search centroids lie far to the west of their vFs (compare the CDFs in SF2a). On the rhumb-line-only hypothesis, terrain seen while flying has no implications for where they should search. It does not, therefore, explain the very large effects of release longitude on where search centroids fall relative to the longitudes of the vFs (see SF1a).

The publicly available animations (movies) of the fix sequences of each recruit (<u>OSF</u>) show that some recruits in each group made a run toward the food, some toward the vF, and some to the RS. They also show that many recruits searched systematically toward more than one target in different phases of their search (see for example, Fig 1c). To capture this aspect of the data, we devised a measure of the strength with which multi-fix segments of a search gravitate to a target location (Fig. 6).

With reference to Fig. 6, the efficiency with which the course segment defined by two successive fixes moves the bee closer to g is cos(*b*_e_)*r*_i_. The overall progress toward g may be measured by the ratio between the initial distance to the goal (*d*_1_) and the distance as of the most recent fix (*d*_3_). G is the product of these two measures of goal progression: *G*_*i*_ = (*d*_1_/*d*_i_)•cumsum(cos(*b*_i_)•*r*_i_), where *i* indexes over successive fixes in the search, *b*_*i*_ and *r*_*i*_ denote the *b*_e_’s and *r*’s in Fig. 6 (segment bearing errors and distances traversed), and *d*_1_ is the distance to F from the first fix in a search.

The gravitation analysis was first applied to the search flights of hive released recruits and affirms that the searches of hive-released recruits usually gravitate toward F (SF 3).

Fig. 7 shows search fixes from displaced bees and the corresponding plots of gravitations toward F (red curves), vF (blue curves) and RS (cyan curves). Because the range of G is huge (note the differences in y-axis ranges in SF3), the *G*_g_’s have been logged—after converting their negative elements to the reciprocals of their absolute values. The reciprocals of the absolute values of the negative elements are <1; thus, their logarithms are negative. Only the positive values of *G*_g_ indicate the strengths and durations of the runs toward the different targets.

The G measure enables us to parse fix sequences into runs directed at the different plausible targets (F, vF and Rs). A sequence of fixes was scored as a gravitation toward one of these goals when the G score was >50 over the fixes in that segment, *and* greater than the G scores of the two alternative goals. (For overall G statistics, see SF4-SF6.)

More than half the bees in every experimental group gravitated one or more times toward the true location of the food at some point in their search (blue bars in Fig. 8). This fraction was significantly greater than the equivalent fraction for the vF and RS goal locations (red and orange bars). This fraction was 0 for one of the control groups taken from a distant hive (R5c); for the other (R2c), it was 0.38. The fraction of hive-released bees showing strong gravitation to F was 0.82. This fraction was approximated by three of the groups released at displaced sites in familiar territory (R1, R5 and R6). In short, displaced recruits released in familiar territory were almost as likely to make at least one run toward F in the course of their search as were recruits released at the hive, while recruits unfamiliar with the release terrain rarely made such runs.

**Fig. 8.**
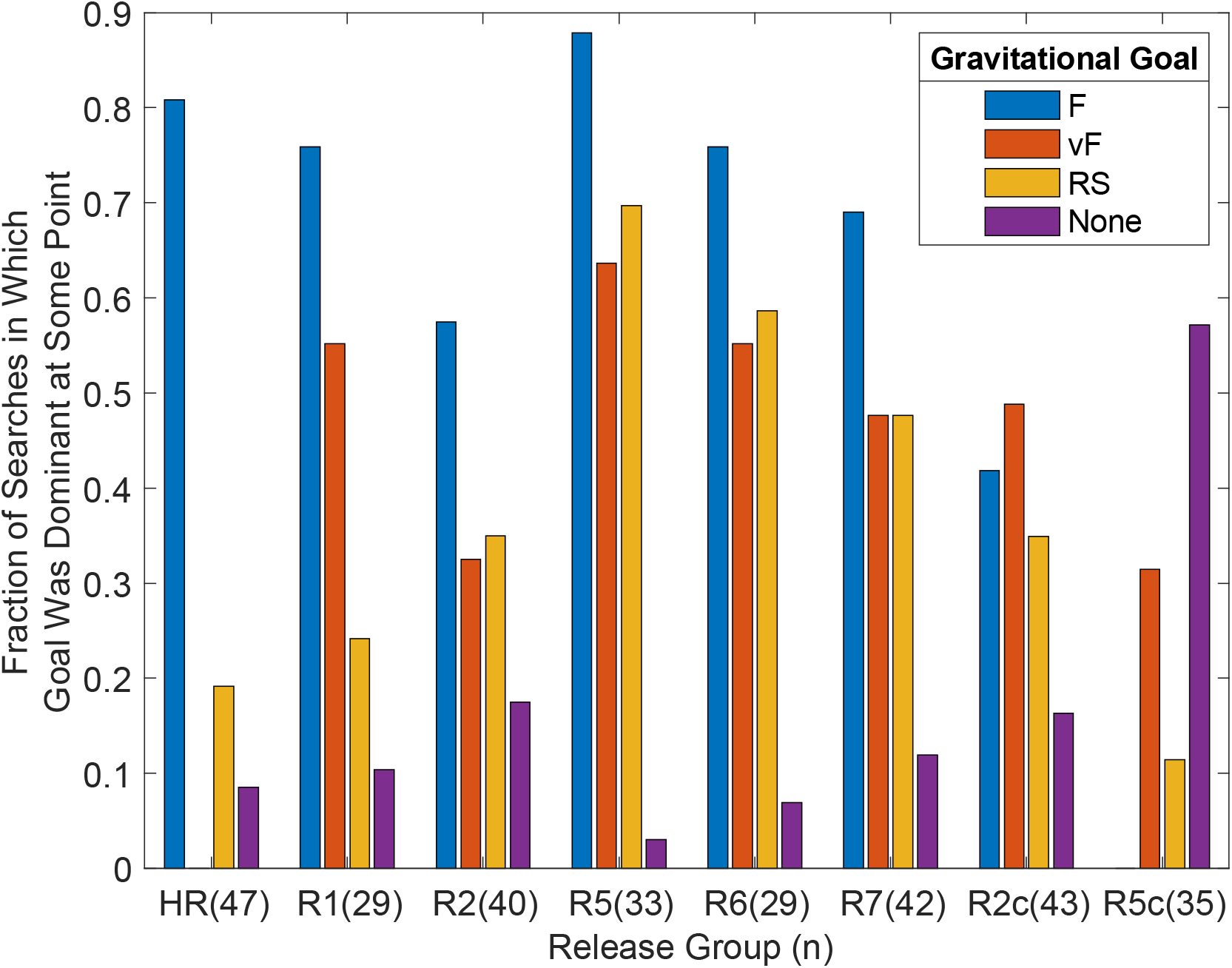
The fraction of the searches in each release group that included at least one strong gravitation (G>=50) toward the goal indicated by the color of the bar (blue for F, red for vF, orange for RS). Purple bars give fraction whose search had no dominant segments. The numbers in parentheses are the n’s. X^2^ tests for differences in the F:vF (blue: red) proportions and F:RS (blue: yellow) proportions across groups R1,R2,R5,R6&R7 yield p’s<< .0001. R2c and R5c give the results for the two control groups. A blue bar is absent from R5c because the fraction directed to F = 0; red bar is absent from H, because vF ⍰ F.

In the animated fix-by-fix sequences (movies viewable at <u>OSF</u>), one sees recruits approach F from diverse and distant locations in every compass direction. These approaches are examples of the distinguishing feature of navigation based on a Euclidean map—the ability to set a course to any location within the map’s frame of reference from any other location within that frame, whether or not the navigator has pre-existing locale information (images) for either location. In assessing the force of the evidence from our experiments, there is no substitute for viewing the individual animated search sequences. The 2-5 most impressive sequences in each release group have been collected into a single movie, *GreatestHits.mpg*. Like the complete sets of individual movies for each release site, it can be viewed by going to this <u>OSF</u> site and clicking on the file. It should open in the reader’s browser in whatever application the browser summons to display .mpg files. We suggest also reading the *Introduction to the Search Movies.docx*, at the same site.

## Discussion

The current results may be understood on the hypothesis that the bee brain, like the vertebrate brain, constructs a Euclidean cognitive map of the terrain in which the animal forages. A Euclidean map is a vector space on whose vectors distances and directions are defined (that is, computable from the location vectors) and Euclid’s parallel postulate holds. For simplicity and economy of representation and computation, the basis vectors are commonly taken to be orthogonal. Navigational computations—vector addition, vector inversion, Cartesian-to-polar and polar-to-Cartesian conversion—operate on the vectors in this space or on displacement vectors derived from its location vectors. Among the location vectors in this space is one that marks the animal’s current location; it is updated by path integration. Thus, the navigator knows *approximately* where it is on its map, although, for displaced recruits, that knowledge is further corrupted by inaccuracies in their estimation of the location of their release site.

Location vectors also give access to terrain views and to goal-properties that may prove useful in the future, such as the color, odor and shape of the flowers the foraging bee has visited (4,6, 18, 19, 20 Chaps 14 & 15). In mammals, non-spatial information accessed by way of location vectors is represented in other vector spaces (21-23). Vector representations of non-spatial features (e.g. odors) are also seen in insects (24-26). Vector representations of diverse aspects of the experienced world appear to be neurobiologically common across disparate phyla.

For decades, Tolman’s hypothesis that rodents form a map of their environment on which they base their navigation (27) was widely disdained (28). In recent decades, however, partly in the light of extensive electrophysiological evidence (29), and partly from an appreciation of the importance of path integration in animal navigation (20), the hypothesis has been widely embraced by cognitive scientists and vertebrate neuroscientists (30-33)—but not by the insect navigation community (34-37). The history of the cognitive map concept in mammals illustrates nicely that a final settling of the debate did not come from behavioral experiments only but from combined neurophysiological work.

That the metric map hypothesis explains our results does not, of course, imply that it is true. It will not prove true if and when another hypothesis is proposed that explains our results without positing a metric space containing location vectors and that is confirmed by neuroscientific results. A reviewer has suggested the alternative portrayed by the bidirectional arrows in Figure 9. On the reviewer’s hypothesis, the forager has stored a large number of bidirectional rhumb lines between the hive and various locations in the environment from which it has made homing flights. The hypothesis further asserts that the dance “primes” (summons up) the inverted rhumb line associated with the F location and the terrain views around it and the panoramas seen in flying toward it from the hive.

**Figure 9.**
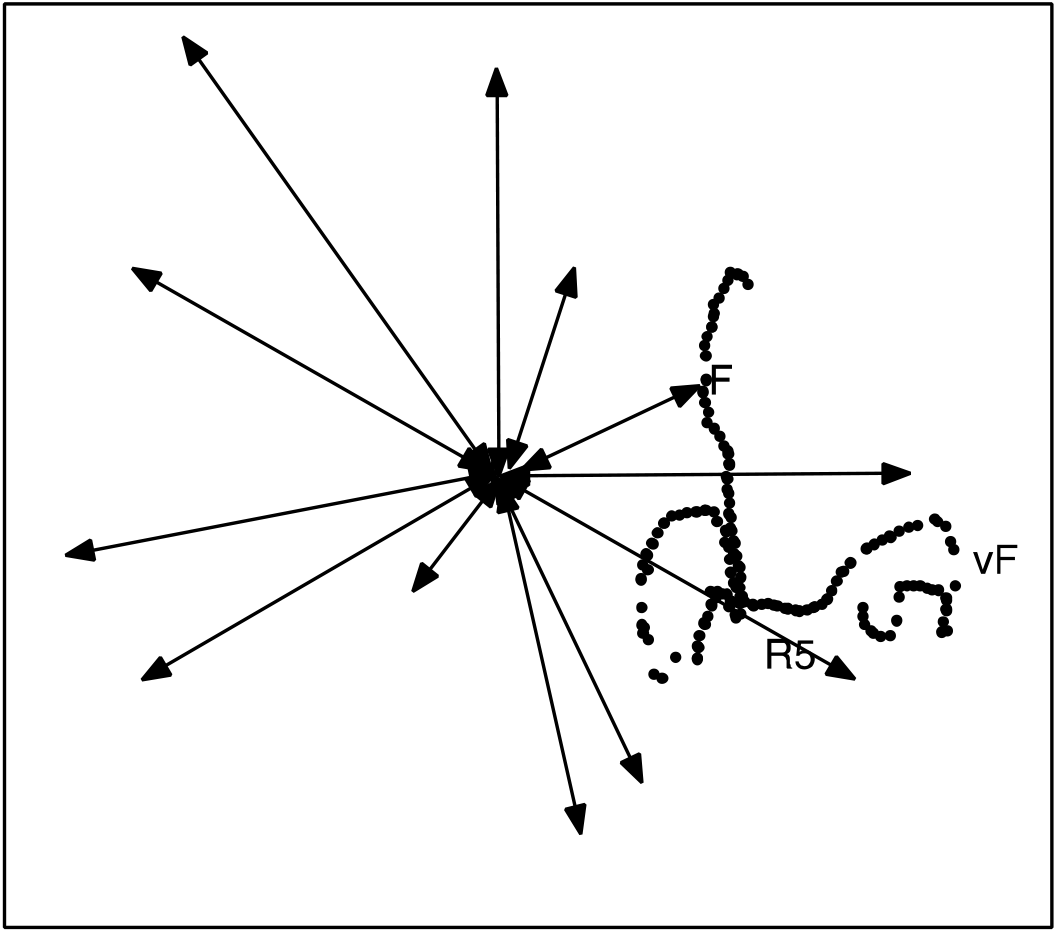
The bidirectional arrows indicate invertible rhumb lines between the hive and landmarks from which a forager has notionally homed. A reviewer-proposed alternative to the map is that our results may be explained on the hypothesis that the dance primes the rhumb line from the hive to F. The dots are a replot of the fixes in Figure 1c.

The vectors indicated by the bidirectional arrows in Figure 9 are metric displacement vectors; one element specifies a compass direction and the other a distance. They are bidirectional because the displacements come in invertible pairs, each pair consisting of the rhumb line from a location to the hive and the rhumb line from the hive back to that location. The reviewer’s hypothesis explains why the vector flights of the displaced foragers are perturbed, because they do not see the primed terrain and panorama. However, it does not explain why the perturbations are release-site specific (Figs 2, 3 and SF2), because the displaced foragers fail to see the expected terrain and panorama regardless of where they are released. The release-site specificity implies that the vector flights are perturbed not only by what the forager *fails* to see but also by what she *does* see. The reviewer’s hypothesis explains the former but not the latter.

The reviewer’s alternative hypothesis also fails to explain the many segments of the ensuing searches that gravitate strongly toward F along compass bearings that are not those of any of the bidirectional arrows in Figure 9. On the alternative hypothesis, foragers can fly from a non-hive location toward F only if F happens to fall on a line from the hive through the start of the F-oriented flight segment. The fixes replotted from Figure 1c into Figure 9 are one of many examples in our data of search segments strongly gravitating toward F along lines turned far away from the line that passes through both the hive and the start of the segment. In Figure 7b, one sees a segment directed to vF, followed by a segment directed from vF straight to F, followed by a segment directed from F straight back to vF, followed by a second segment directed from vF straight back to F. The three latter segments are inexplicable on the alternative hypothesis. By viewing the movies of all the searches, one may confirm that a majority of the strong gravitations to F are similarly inexplicable on the proposed alternative hypothesis.

One may also wonder why on the map hypothesis all the recruits do not fly directly from vector termination to F. The answer is that the courses set by displaced recruites are subject to multiple sources of error that together often make it impossible to set a course that brings the navigator close to the intended goal. The dance itself contains substantial error because the rhumb line and the location vector for the source from which the dancer returned both depended on path integration, which has a cumulative error that depends on the linear distance traversed in reaching the source (38, 39). There must be further error in the interpretation of the dance by the forager that followed it, because the interpretation depends on gravitational information about deviation from the vertical (for the bearing of the rhumb line) and counts of the number of waggles or the durations of the waggle runs (for the distance). Thirdly, the mapped locations of the landmarks and the terrain views also depend on dead reckoning, so there are substantial errors in the map itself (<u>see, for example, the Waldseemüller map</u>)—both the dancer’s map and the recruits’ map. Uncertainty about where they began their vector flight further augments a displaced recruit’s dead reckoning error. Finally, determining where one is on a map from sightings on landmarks is itself extremely error prone (40-42). In short, setting an accurate course from where one is to a goal depends on the accuracy with which both locations are known. Large errors in these locations lead to large errors in the courses set, hence an inability to find one’s goal.

The inadequacy of the reviewer-proposed alternative does not, of course, preclude other hypotheses that may be advanced in the future. Be that as it may, our results lend credibility to the map hypothesis for insects, because currently published alternatives (35) do not explain them and the map hypothesis does explain them.

## Methods

### Experimental site, honeybee colony, experimental design

The experiments were carried out in 2015, 2016 and 2017. The experimental site was a structured flat agricultural landscape with grass fields stretching to the east of the area scanned by a radar (located at R/H, Figure 1, coordinates: 50°48’52.21”N, 8°52’20.43”E) with trees and bushes, pathways, and creeks close to the Großseelheim village (Germany). The whole grass area east of R/H was frequently cut by the farmers. The experiments started early August at a time when natural food sources appeared only west of R/H in gardens of the village (marked bees from our colony were frequently seen by the garden owners) and a few small spots along the creeks. Scattered flowers of *Lapsana communis*, a yellow blooming composite providing only pollen, appeared 3 –4 days after the last cut rather equally distributed over the whole grass area and disappeared with the next mowing soon after. No denser patches of these flowers were ever seen at the area around F.

The bee colony in an observation hive with about 3000 bees was introduced to the area 10 to 14 days before the experiments started in order to familiarize the foragers with the area. During this period and throughout the experiments the bees appearing at the hive entrance were marked individually with number tags. The two digit number tags had 5 different colors and were glued to the thorax of the bees in 4 different directions relative to the body length axis allowing to mark close to 2000 different bees (numbers sensitive to the direction of reading were excluded, eg. 66 and 99, 69, 96). All bees used in the experiment as trained bees (dancers) or recruited bees (recruits) were individually marked and full protocols were established. The trained bees were additionally marked with a white dot on the abdomen including the period of training from the hive to the final feeder location. No bee used as trained bee at any stage of training was used as a recruit. Dances of bees for natural food sources were observed throughout the experimental period, and no dances indicating a source to the East (the radar scanned area) were observed, except by those trained to dance to F.

Recruits were caught at the hive entrance after they had followed a dance of an experimental dancer using a transparent marking device (a tube with a stopper) and transported to the release site in a dark box. The time interval between catch and release was close to the same for all release sites including the control experiment. Returning recruits were removed from the colony when they arrived at the hive entrance and after the radar transponder was collected. The feeding place F for the trained bees was located at a distance of 397 m and a direction of 71° east of N. The trained bees served as the dancers. The feeder was an unscented plastic container standing on a small table. Recruits rarely found the table and landed at the feeder (3 out of 47 hive released bees, none of the displaced recruits), most likely due to the unnatural conditions and the lack of odor. Bees without abdomen marker were frequently seen dancing for natural food sources in the west of the colony. Recruits were released either at the hive (R/H) or at one of the five additional release sites (R1, R2, R5, R6, R7, Fig. 1 a).

### Tracking by harmonic radar and segmentation of flight trajectories

We used a system with a sending unit consisting of a 9.4 GHz radar transceiver (Raytheon Marine GmbH, Kiel, NSC 2525/7 XU) combined with a parabolic antenna providing approximately 44 dBi. The transponder fixed to the thorax of the bee consisted of a dipole antenna with a low barrier Schottky diode HSCH-5340 of centered inductivity. The second harmonic component of the signal (18.8 GHz) was the target for the radar. The receiving unit consisted of an 18.8 GHz parabolic antenna, with a low-noise pre-amplifier directly coupled to a mixer (18.8 GHz oscillator) and a downstream amplifier with a 90 MHz ZF-filter. A 60 MHz ZF-signal was used for signal recognition, leading to a fixing of the bee carrying the transponder.

The transponder had a weight of 10.5 mg and a length of 11 mm. We used a silver or gold wire with a diameter of 0.33 mm and a loop inductance of 1.3 nH. The range of the harmonic radar was set to 0.5 nautical miles. The frequency of radar fixes was every 3 s. The raw radar output was captured from the screen at a frequency of 1 Hz, stored as bitmap files, further analyzed offline by a custom-made program that detected and tracked radar signals (fixes), and converted circular coordinates into Cartesian coordinates taking into account multiple calibration posts in the environment. Finally, the fixes were displayed in a calibrated geographic map created with the software Pix4D from aerial images (43) taken with a commercial drone (DJI Inspire). In the rare cases when no fixes were received from a bee for more than 30 s, the flight trajectory was interrupted, and the last, as well as the first, fix before and after interruption were marked.

#### Analyses of flight trajectories

Segmentation of flight trajectories: A total of 369 recruits were tested. Typically, recruits performed a sequence of three sequential flight segments: the outbound vector flight; the search flight; and the inbound homing flight (Fig. 1 a). Homing flights were not further considered here because all bees returned home on fast and straight flights. Some of the recruits released at R1 or R2 initiated a straight homing flight after close exploration around the release site without performing a vector and or search flight. These recruits were not included in our analyses. The transitions from the rather straight vector flight to the search flight and the search flight to the homing flight were characterized by a sharp turn of ≥60° with straight stretches before and after the turn with at least three fixes each (Fig. 10).

**Fig. 10.**
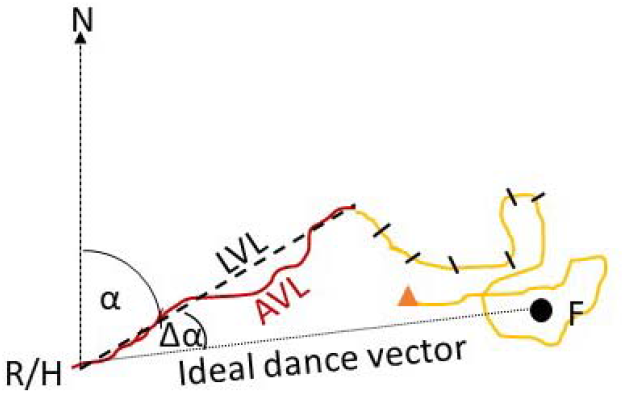
Parsing of a recruit’s radar fixes into an outward-bound vector, a search component, and a hive-bound vector was done by an algorithm. Magenta = vector portion; yellow = search portion; cross-ticks on search portion indicate fixes at 3s intervals; red triangle marks beginning of hive-return vector. Bearing α: bearing of terminal vector fix in degrees clockwise from N. Length: line between release site and terminal of vector flight (dotted line). Speed: interval between start and end of vector flight and the real distance flown (AVL: accumulated distances of fix-fix segments). Straightness: proportion of LVL and AVL. F = food location.

### Statistics

Statistical analyses were run by scripts in R or in Matlab(tm). The code that implements the G computation has been uploaded <u>here</u>, along with the raw data, all of the results of our analyses, and the code that produced those results. A one-way ANOVA was used to compare vector parameter data from displaced groups to the data from the hive-released group. The proportions in Figure 8 were compared by chi square tests.

## Acknowledgments

We are grateful for expert assistance during the experiments by Z. Hu, Y. Qu and K. Tan. The honeybee colonies were kindly provided by Dr. R. Büchler, Bieneninstitut Kirchhain, Hessen. We are also grateful to the farmer, Mr. Lemmer, who gave us permission to work in his agricultural fields. Our thanks go to Dr. R.R. Fitak for his advice on circular statistics and bimodal models. We are also grateful to Dr. G. Galizia for giving us most helpful feedback on an earlier version of the manuscript. Julian Petrasch and Dr. Tim Landgraf provided us with the aerial view of the landscape.

## Data and Code Availability

**The raw data and the code by which they were analyzed and the figures produced may be accessed at https://osf.io/a59rs/files/?view_only=6da39e230c8d4072b27ab70ccecb06e2. The raw data for every figure may be found at this link**.

## Related manuscripts

none

## Supplementary information

Link to Video

### Control experiment

In order to test whether recruits behave differently if they had not explored the test area during their orientation flights, we located a colony in an area 4400 m SE of the test area behind a hill (50°47‘35.78‘‘N, 8°55‘36.32‘‘E). Both the ground structure and the skyline differed substantially from the conditions in the test area. The dancers were trained in an urban area to a feeder 373 m away from the hive under an angle of 69° (50°47‘40.87‘‘N, 8°55‘53.17‘‘E). The flight distance and direction of the dancers resembled the conditions in the main experiment. The recruits were caught at the hive exit in the same way as in the main experiment, stored in a catching device and transported within 20 minutes to one of two RSs (R2, R5) in the test area, equipped with a transponder and radar tracked. It might be thought that the fact that a troop of bees had foraged at F created lingering odor cues there that attracted the displaced recruits even from great distances and diverse directions. To control for that possibility, we installed a second hive at the R/H location and trained a troop from it to forage at the table at F during the time the control bees were released at R2 and R5.

Recruits of the control experiments performed much less vector flights than recruits of the main experiment: At R2: 48 % (n=23) of the control bees as compared to 70% (n=34) of the bees in the main experiment, at R5: 13% (n=30) as compared to 79% (n=34). Flight distances of vector flights in control experiment were significantly shorter: R2: average accumulated vector distance: 159±69 m, average linear vector distance: 128±57 m) as compared to those in the main experiment (average accumulated distance: 366±156m, average linear vector distance: 314±133 m (p<0.01 for accumulated distance, p<0.05 for linear distance); R5: average accumulated distance: 158 ±80 m, average linear vector distance: 128 ±46m as compared to those in the main experiment: average accumulated distance: 274±149 m, average linear vector distance: 223±127m, Man-Whitney U-Test p<0.01 for accumulated distance, p<0.05 for linear vector distance). Flight speed was significantly lower in control animals (R2 control experiment: 3.5 ±0.86 m/s, R2 main experiment: 6.7±1.7 m/s, Man-Whitney U-test p<0.01). Similar results were found for the release site R5. The directions of vector flights were significantly different in R2 released bees as expressed in the deviation from the ideal vector Δα (control experiment (Δα=34.3±14.7°. main experiment: average Δα= -7.5±16.4°). Thus the vector flights in the main experiment were more tilted to F. This difference is highly significant (p<0.01 Man-Whitney U-test). No significant difference was found for R5 released bees.

Thus, recruits in the control experiment performed significantly less vector flights at both release sites and shorter vector flights. This difference was larger at R5 than at R2. The direction of the vector flight was not tilted toward F as it was for the recruits in the main experiment. The distances flown during search flights were significantly different for R5 but not for R2. It appeared that recruits starting at R2 may have found the area less unfamiliar than when starting at R5 since they had explored an urban area with houses, gardens and streets, and the area west of R2 resembled such structures at least in part.

### Vector flights

The initial phase of the recruits‘ trajectory is dominated by the information they received from the dancing bees (vector flight). SF 1a and b document these conditions by plotting the vector flight fixes and the endpoints of the vector flights in relation to H, F and vF.

**SF 1.**
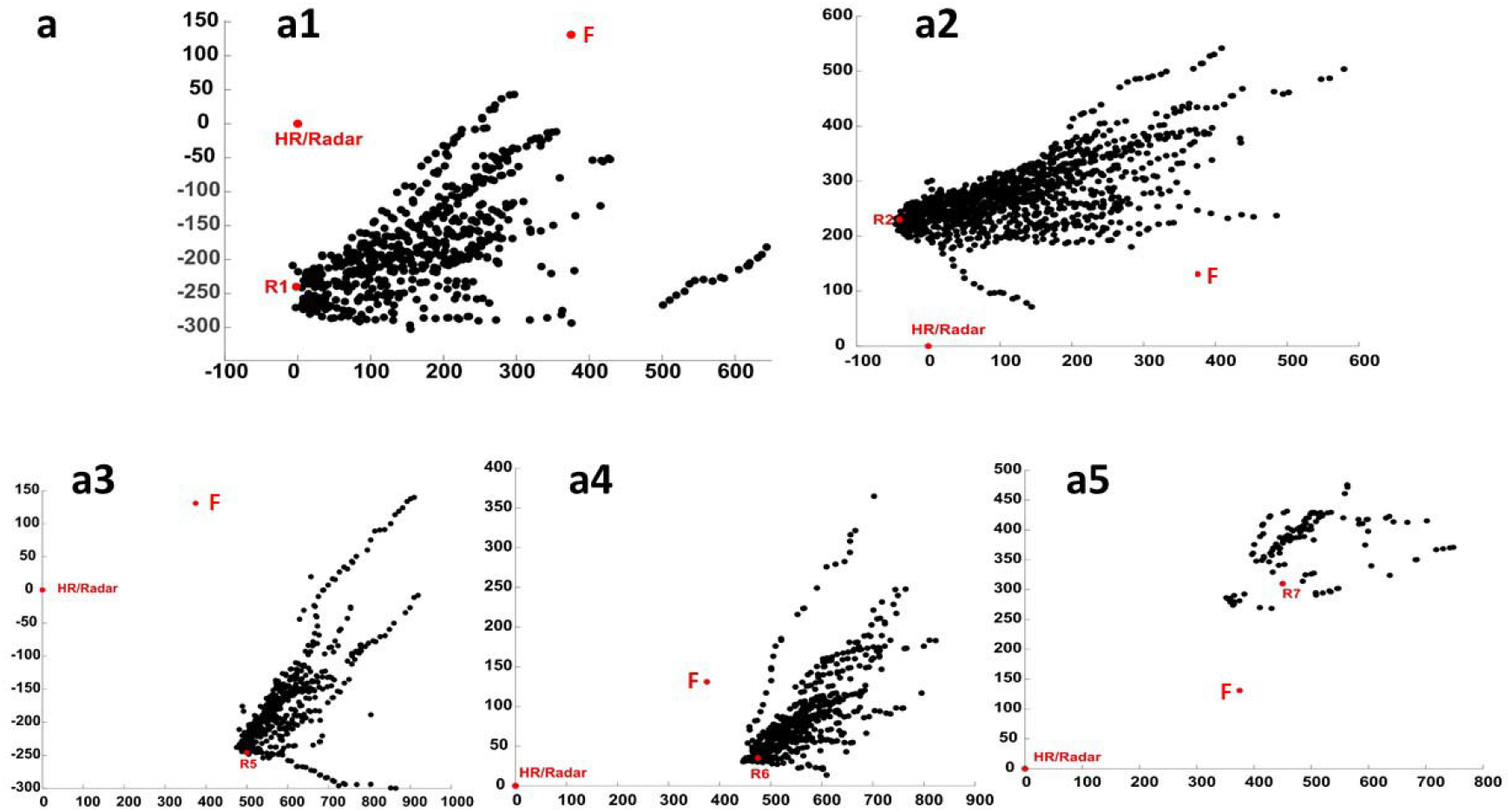
Radar fixes of vector flights for all recruits released at the 5 release sites other than the hive (a1 – a5).

### Search Centroids

The centroid of a search is the mean of the longitudinal and latitudinal coordinates of the fixes (a 2-element vector, the first element giving the average of the east-west elements, the 2nd the average of the north-south elements). SF 2 plots the cumulative distribution functions of the longitudinal (SF 2a) and latitudinal (SF 2b) elements on panels with verticals at the coordinates of the release sites, the true location of the food (F) and its virtual locations (vF).

The groups released at or somewhat west of the hive (HR, R1 & R2) are plotted with solid lines; those released east of the food are plotted with dotted lines (R5, R6, R7). The solid distributions in SF 2b straddle the corresponding vF longitudes (which are near the F longitude). By contrast, the dotted distributions fall far to the left (west) of the corresponding vF longitudes. Recruits released at or slightly west of the hive mostly searched 200-600 meters to the east of their release site. None searched more than a few meters to the west of it. By contrast, substantial fractions of recruits released east of F, focused their search well to the west of their release site (and, therefore far to the west of their vF). As a result of this pronounced difference in the compass sectors of the search foci, the dotted CDFs fall much closer to the solid CDFs than would be expected if recruits had no map and just searched around the termini of their vector flight. Put another way, when they terminated their vector flight, displaced recruits tended to focus their searches in the compass sector in which F lay —to the east in those released west of the food and to the west in those released east of it. Similar but smaller shifts are seen in four of the five latitudinal distributions: The distributions for release sites to the north of the hive (R1, R6 and R7) are shifted to the south of their respective vFs, while the distribution for one of the two release sites south of the hive (R1) is shifted to the north.

**SF 2.**
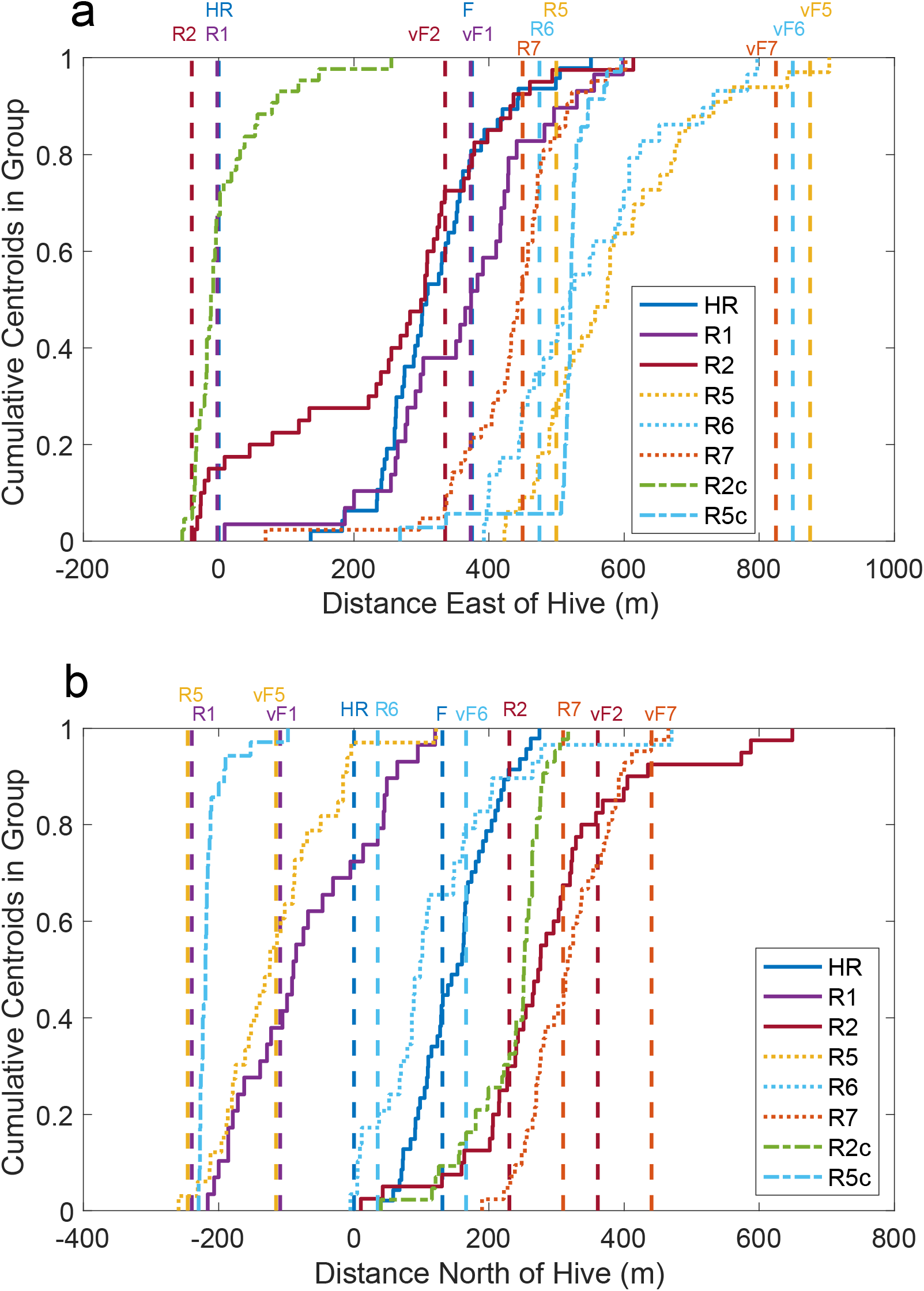
Cumulative distributions of the longitudinal (Panel a) and latitudinal (Panel b) elements of the search centroids. Vertical dashed lines give the longitudes (a) or latitudes (b) of the release sites, the true location of the food (F) and its release-site-dependent vFs in colors that match the corresponding CDF colors. Each little step in a CDF is the centroid for a different recruit in that group.

### Gravitation toward F by hive released recruits

Gravitation toward vF, RS and F were calculated with the formula given in the main text and Fig. 5. The panels in SF 3 plot *G*_F_, gravitation toward F, and *G*_RS_, gravitation toward RS, for 15 of the bees released at the hive. In this group, F and vF are identical, so the vF plot (blue) superposes on the F plot (red), making it invisible in the figure. All but one of these plots shows one or more segments in which the bee gravitates to F—multiple fix-fix segments during which G_F_ rises well above 0. By contrast, the gravitational strength to the release site is generally negative (below the horizontal dashed line).

**SF 3.**
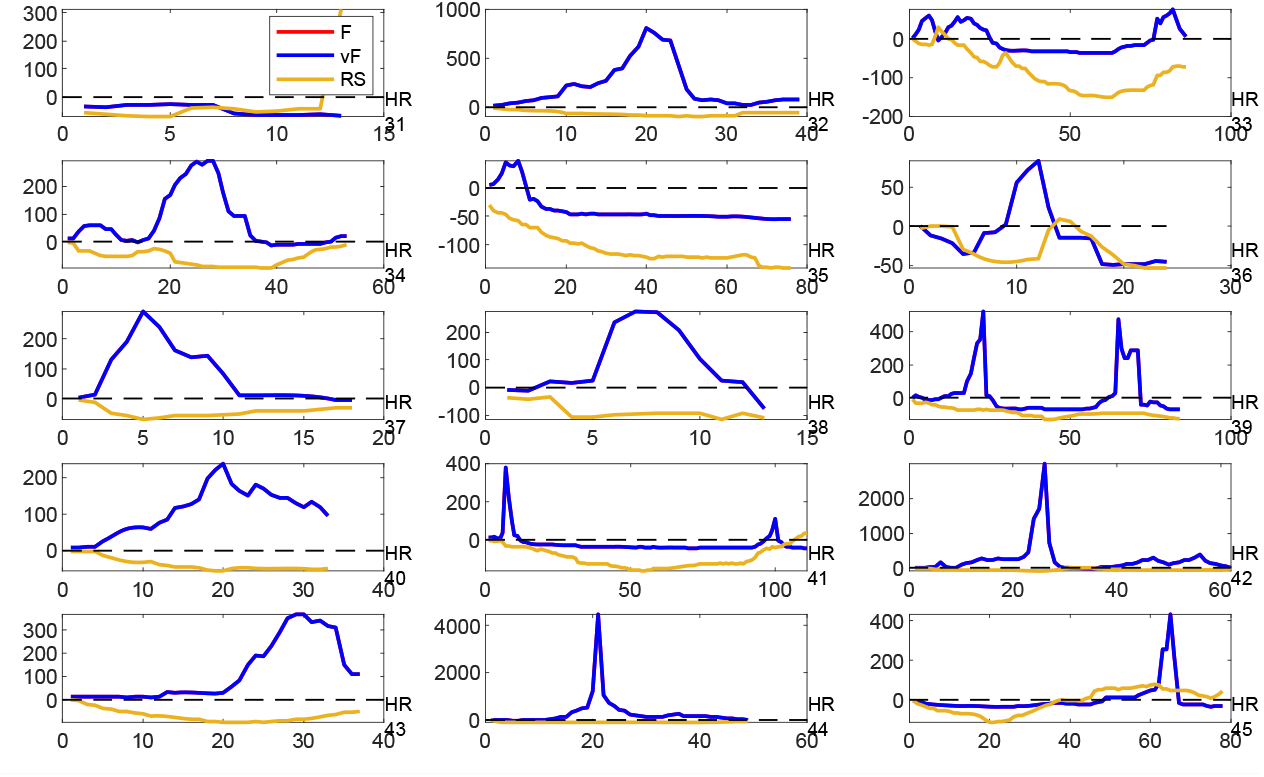
Gravitations toward F (blue) and toward RS (yellow) in 15 hive-released bees. Note the differences in the y-axis scale between panels; the gravitation toward a given goal, G_g_, ranges into the tens of thousands in cases where there is a sustained run in that direction that brings the bee to within a few meters of the goal.

### Gravitation Statistics

**SF 4.**
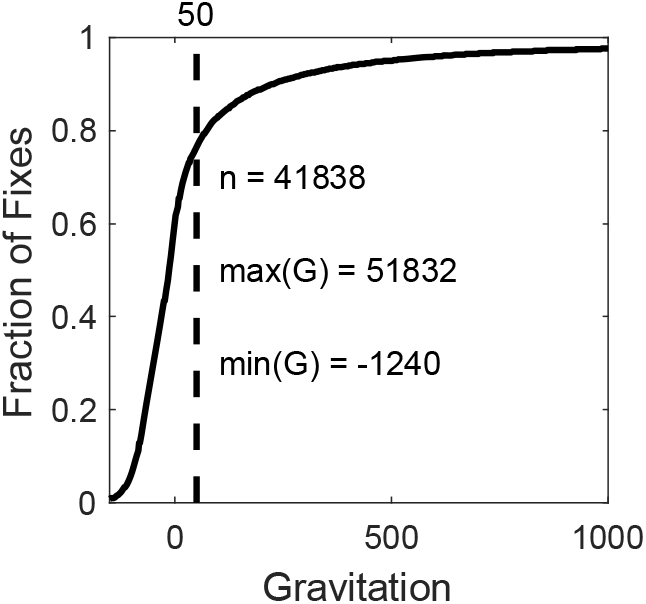
Cumulative distribution of the gravitations scores (all fixes, all conditions). The gravitation algorithm assigns to every search fix beyond the first in the set of fixes from a given recruit gravitation scores (G) for each of three putative goals, F, vF and RS. A sequence of two or more successive fixes with gravitations above 50 (see dashed vertical) are taken to indicate a run toward that goal, provided also that they dominate (are greater than) the gravitations to the other two goals.

**SF 5.**
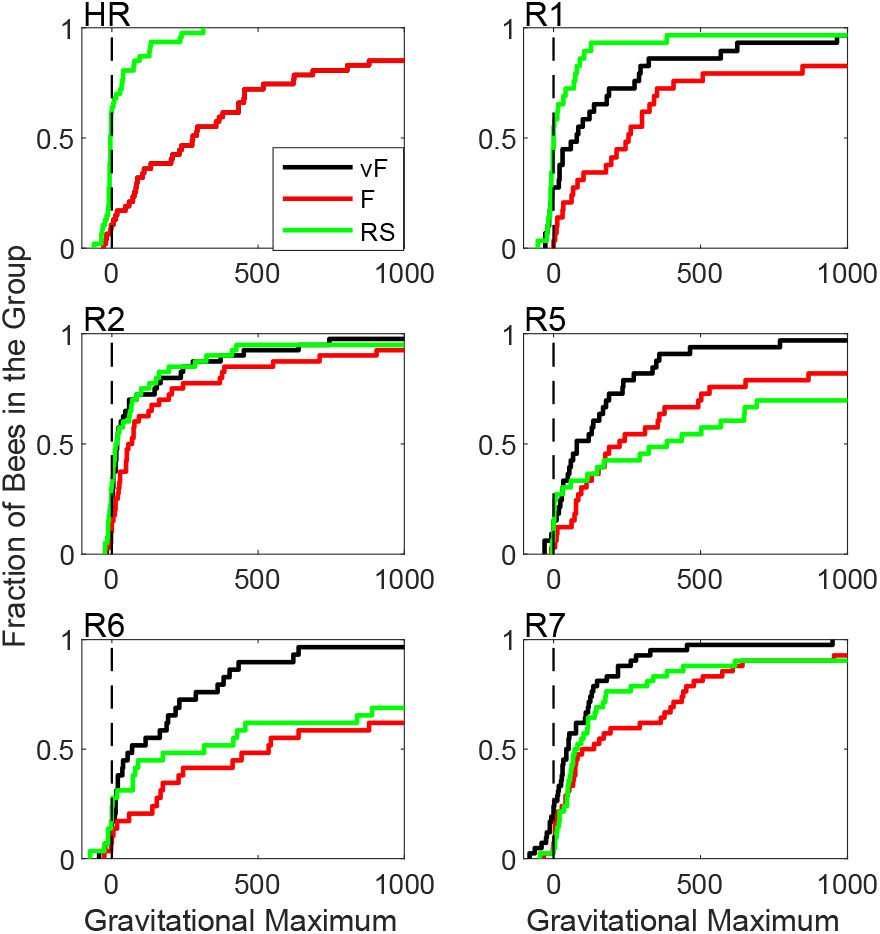
Cumulative distributions of the maxima of the runs directed toward the three different goals (within panel) for the 6 different release sites (between panels). Only recruits from the home hive are included. Portions of the distributions >1000 not shown. Lower cumulative distributions indicate stronger maxima!

**SF 6.**
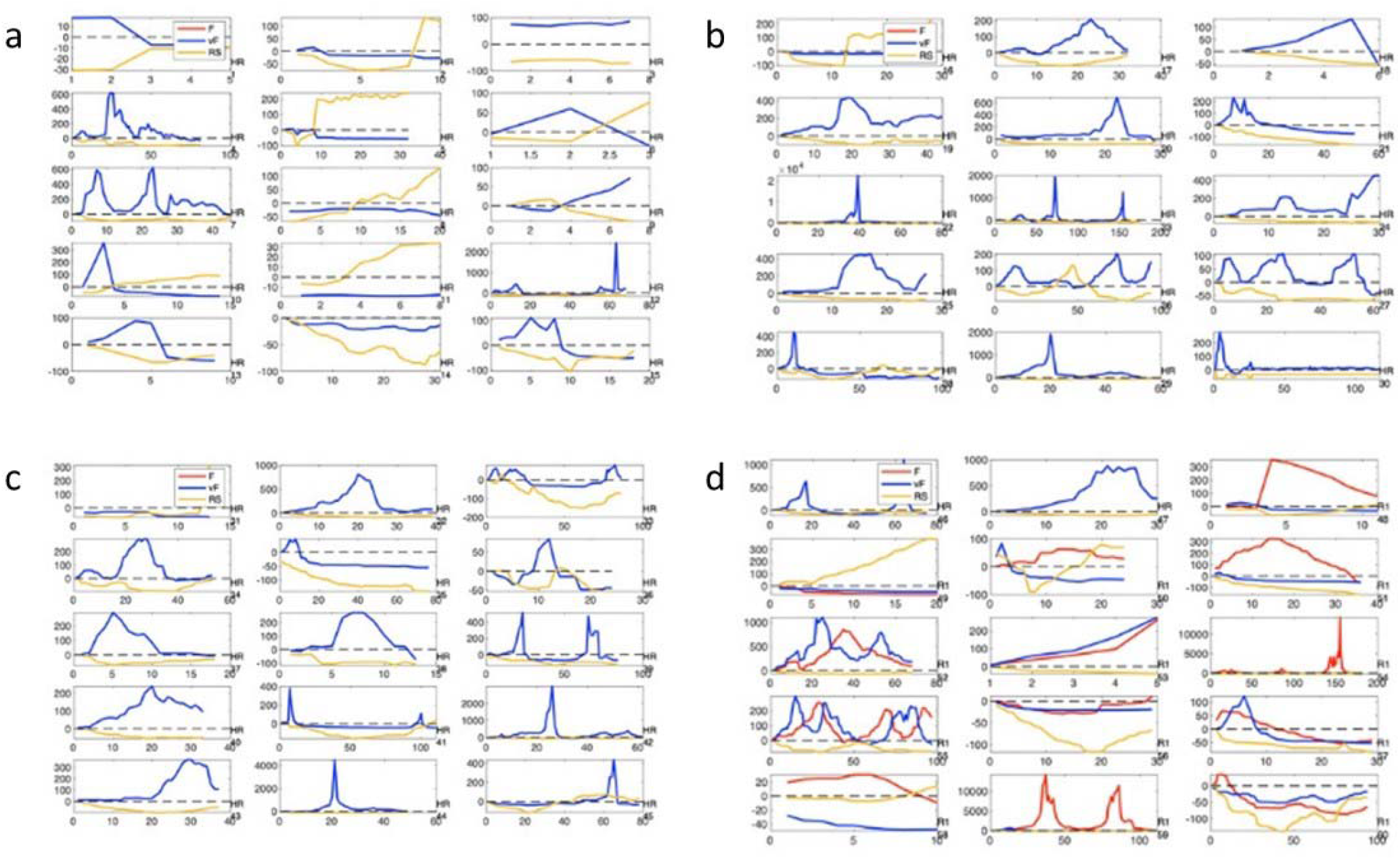
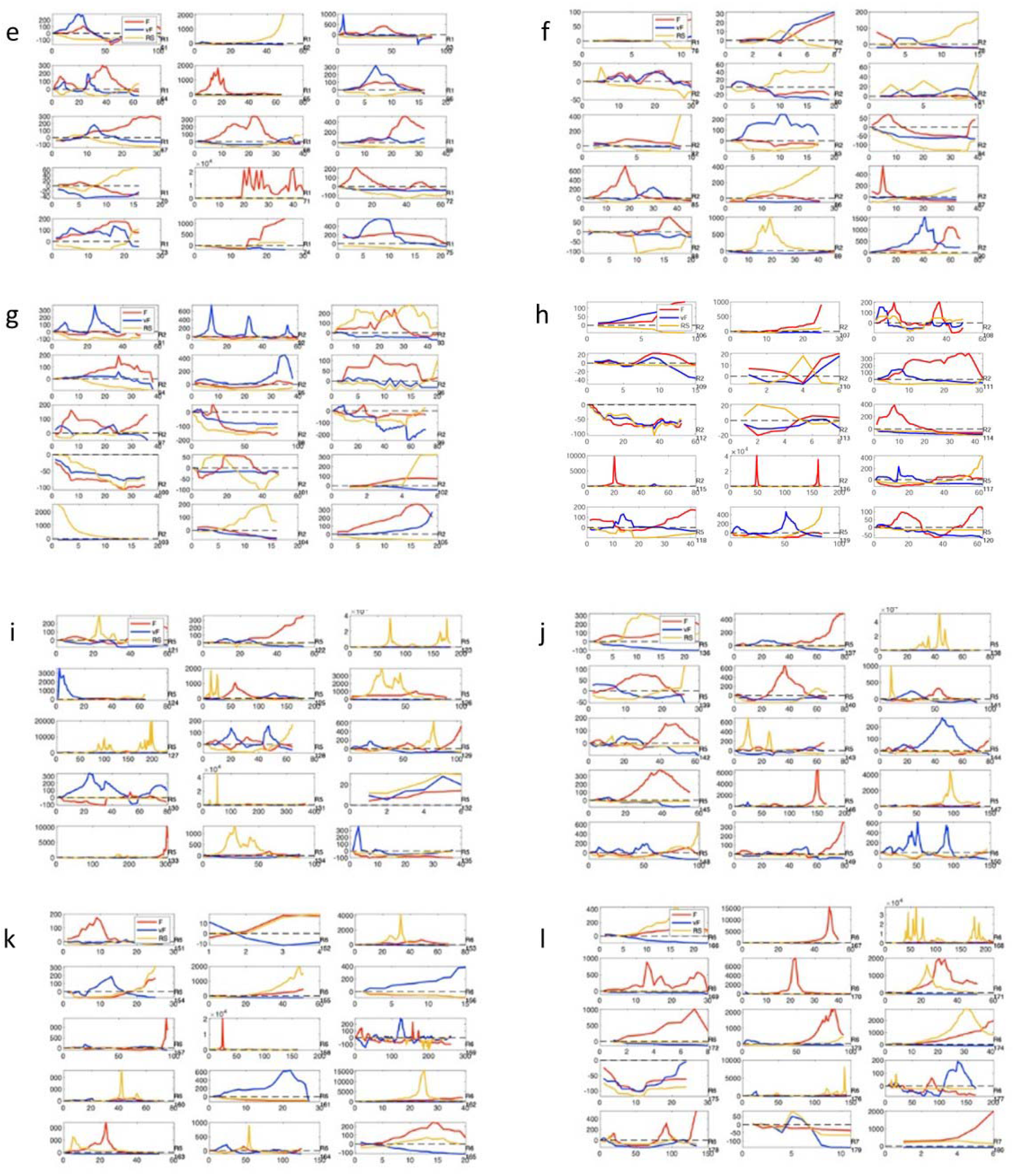
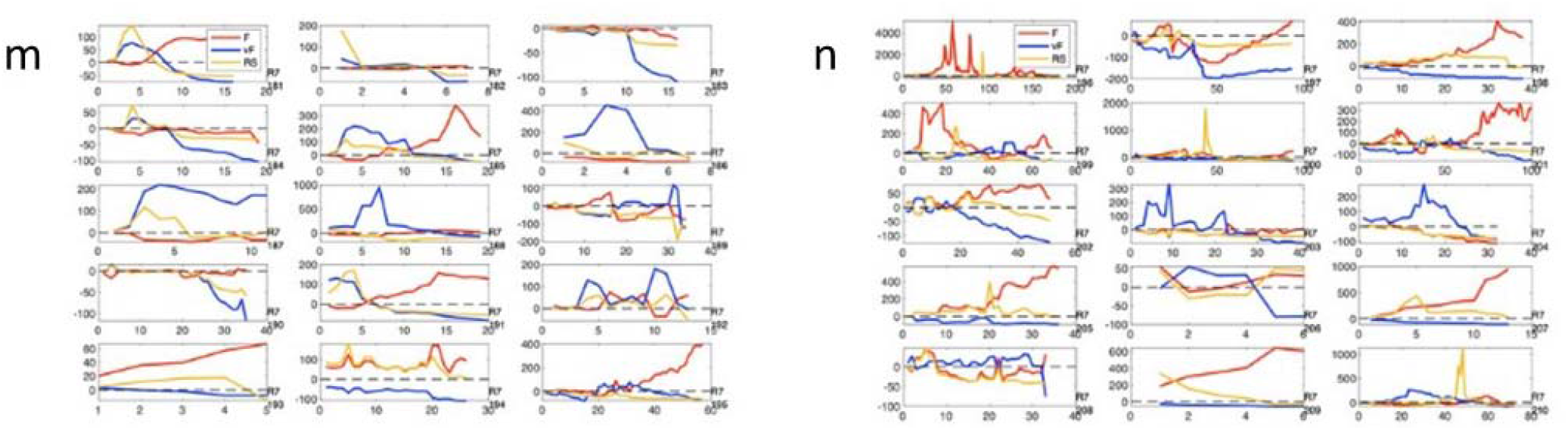
a-n (the above 14 figures, 15 panels/figure). The gravitation plots for all of the searches of recruits from H: G on the y axis and fix number of the x axis. The release site is indicated at the lower right of each panel. G_F_ plotted in red; G_vF_ in blue and G_RS_ in yellow. For the HR bees, vF is identical to F, so the red plot (for F) is obscured by the blue n(for vF). The HR panels do not include those shown in SF-2

### Tables ST-1 to ST-4

The following tables report the means, medians and standard deviations of the vector parameters for each release site and the p values from comparing the displaced distributions to the hive-released distributions using the 2-sample Kolmogorov-Smirnov test. For the *n*’s, see Figure 3 in main text.

**Table ST-1.** Vector Flight Bearing

| Release Site | Mean(°) | Sigma(°) | Median(°) | PvsHR |
| --- | --- | --- | --- | --- |
| HR | 64 | 13 | 67 |  |
| R1 | 72 | 13 | 70 | 0.009 |
| R2 | 78 | 16 | 74 | <<.001 |
| R5 | 53 | 19 | 56 | 0.003 |
| R6 | 62 | 19 | 62 | 0.564 |
| R7 | 38 | 23 | 35 | <<.001 |

**Table ST-2.** Vector Flight Length

| Release Site | Mean(m) | Sigma(m) | Median(m) | PvsHR |
| --- | --- | --- | --- | --- |
| HR | 319 | 132 | 290 |  |
| R1 | 313 | 143 | 317 | 0.841 |
| R2 | 315 | 134 | 314 | 0.852 |
| R5 | 223 | 127 | 197 | 0.002 |
| R6 | 199 | 95 | 174 | <<.001 |
| R7 | 156 | 62 | 144 | <<.001 |

**Table ST-3.** Vector Flight Speed

| <u>Release Site</u> | <u>Mean(m/s)</u> | <u>Sigma(m/s)</u> | <u>Median(m/s)</u> | <u>PvsHR</u> |
| --- | --- | --- | --- | --- |
| HR | 7.64 | 1.94 | 7.75 |  |
| R1 | 6.43 | 2.68 | 6.28 | 0.012 |
| R2 | 5.24 | 2.03 | 5.85 | <<.001 |
| R5 | 3.95 | 4.1 | 2.56 | <<.001 |
| R6 | 4.16 | 2.09 | 4.19 | <<.001 |
| R7 | 6.12 | 3.39 | 7.44 | 0.029 |

**Table ST-4.** Vector Flight Straightness

| <u>Release Site</u> | <u>Mean</u> | <u>Sigma</u> | <u>Median</u> | <u>PvsHR</u> |
| --- | --- | --- | --- | --- |
| HR | 0.94 | 0.05 | 0.95 |  |
| R1 | 0.85 | 0.08 | 0.86 | <<.001 |
| R2 | 0.86 | 0.09 | 0.88 | <<.001 |
| R5 | 0.79 | 0.12 | 0.84 | <<.001 |
| R6 | 0.84 | 0.08 | 0.86 | <<.001 |
| R7 | 0.72 | 0.11 | 0.73 | <<.001 |

