## Supplementary figures and images for "Honey Bees Get Map Coordinates from the Dance"

### Fig. S1

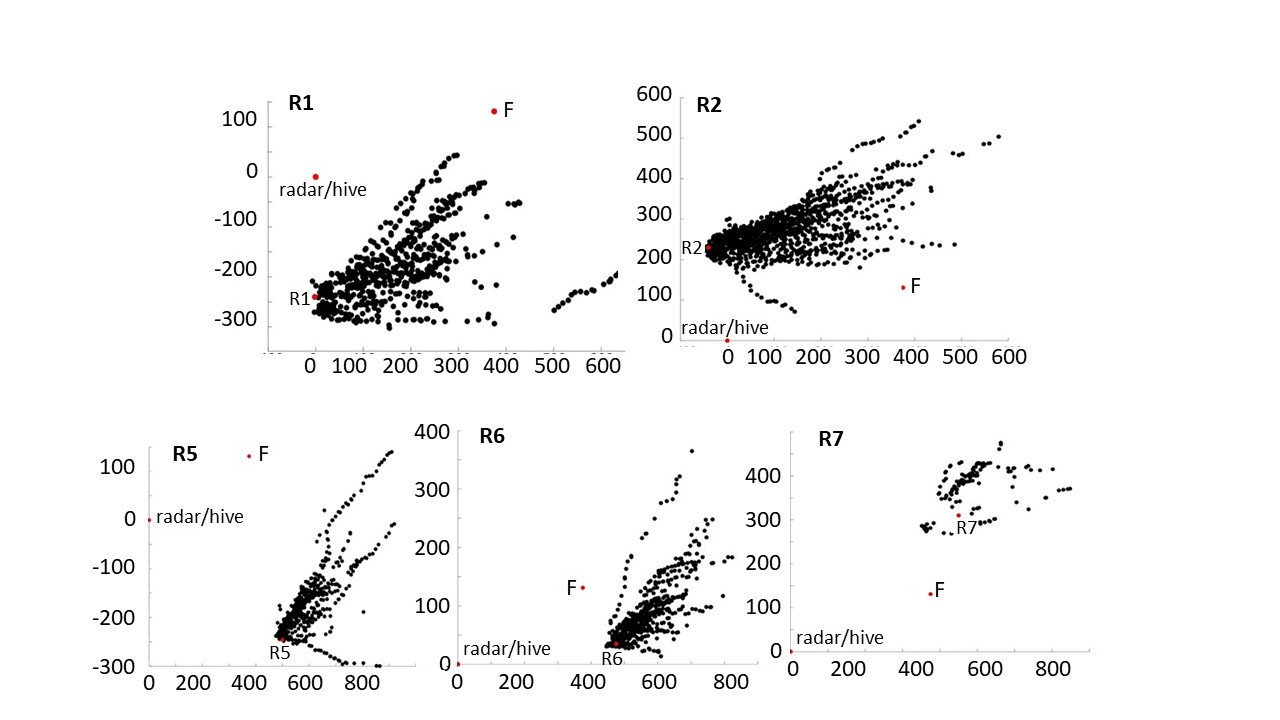

### Fig. S2

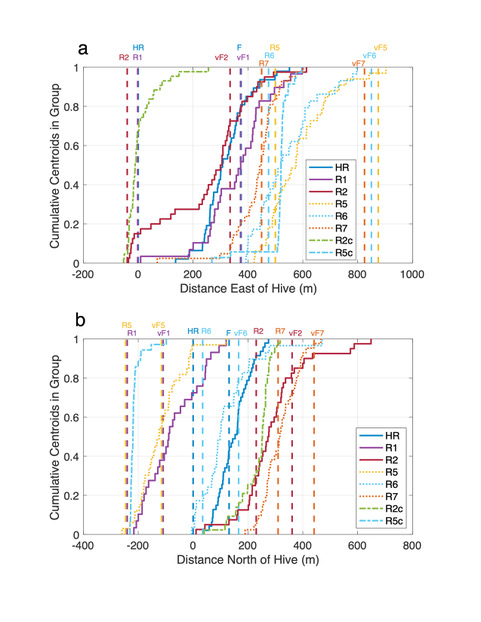

### Fig. S3

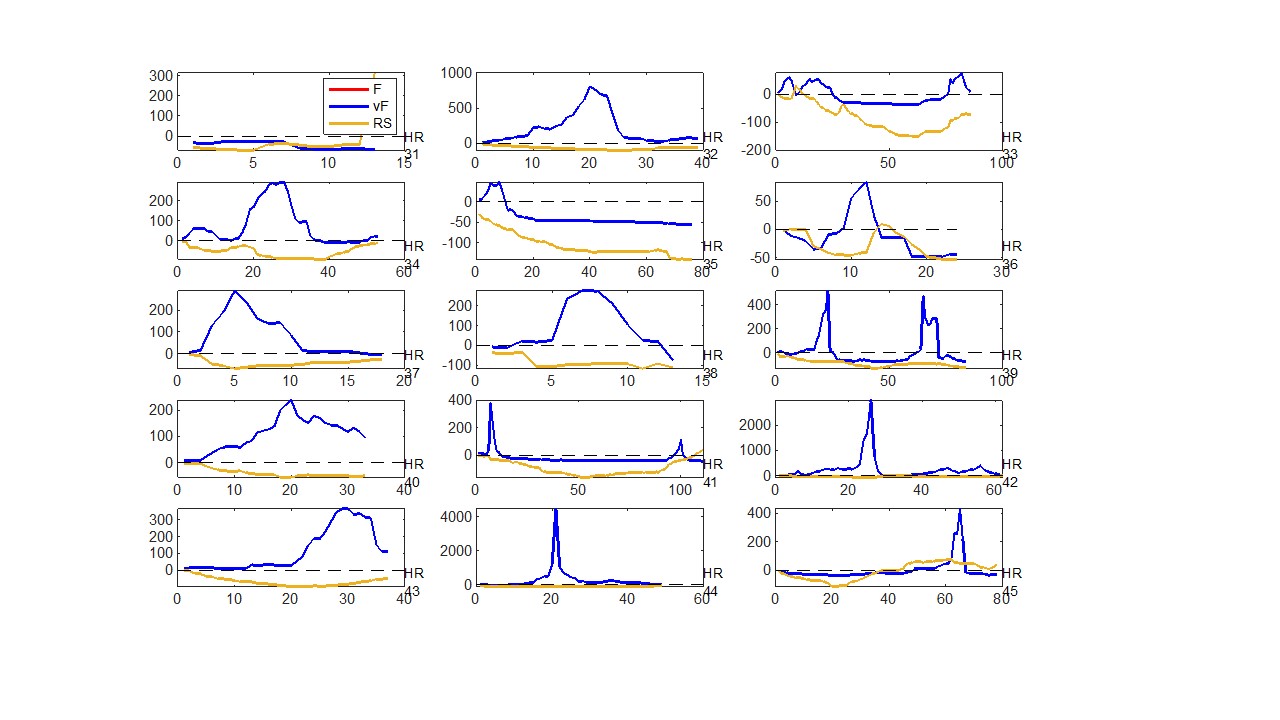

### Fig. S4

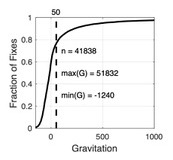

### Fig. S5

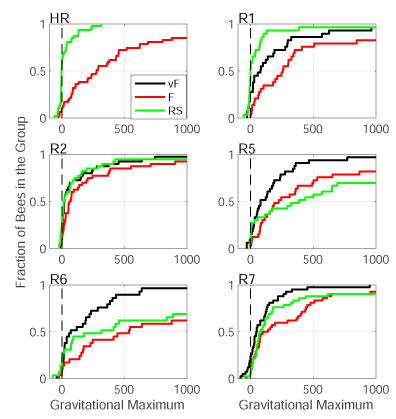

### Fig. S6 i-l

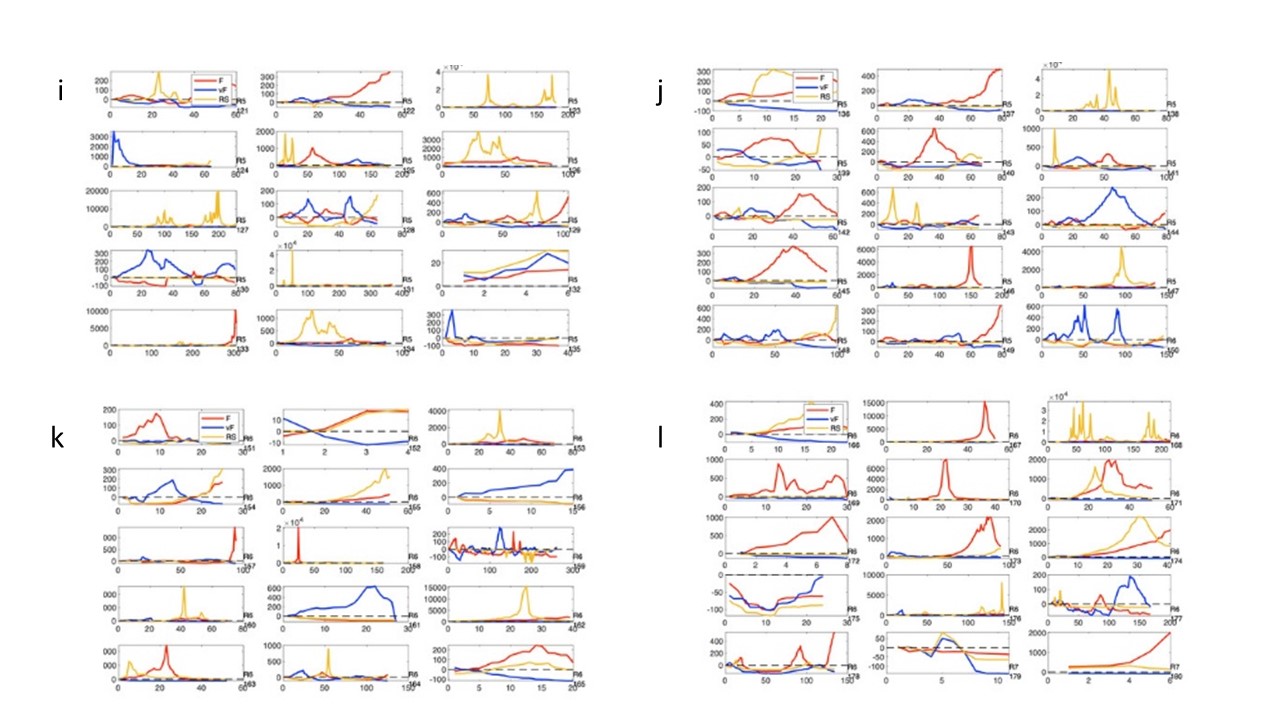

### Fig. S6 m-n

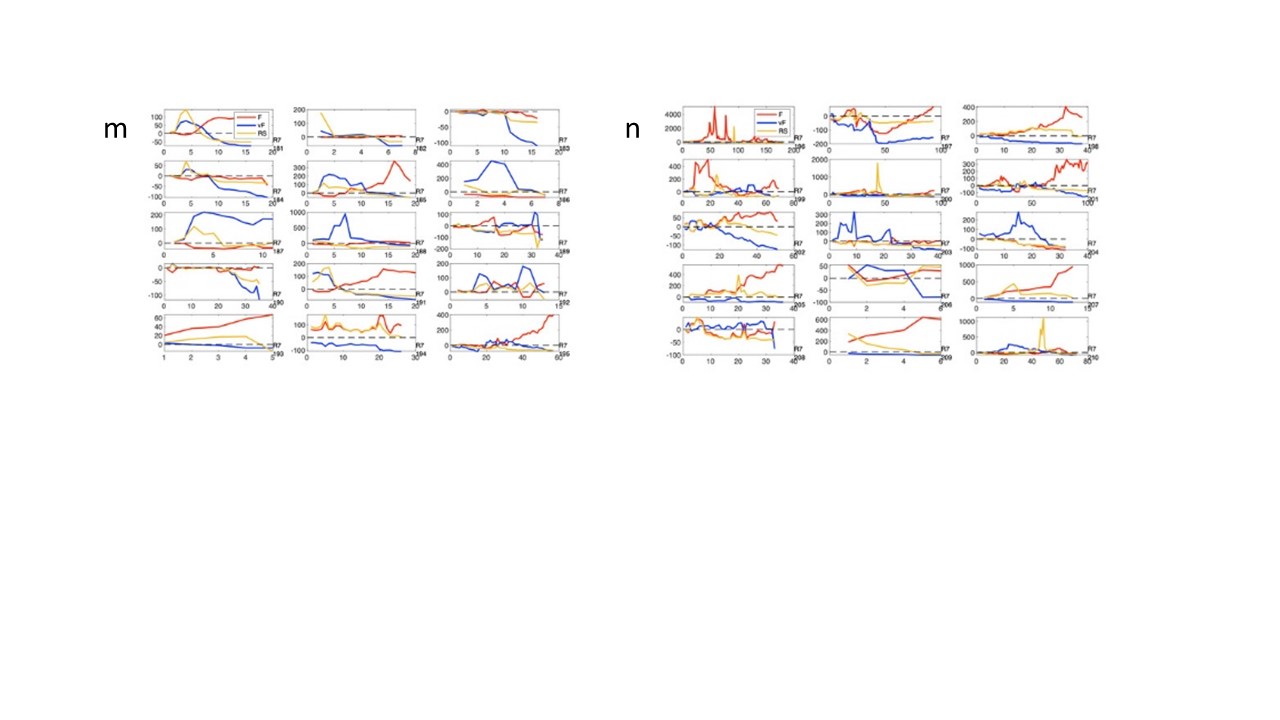

### Fig. S6a-d

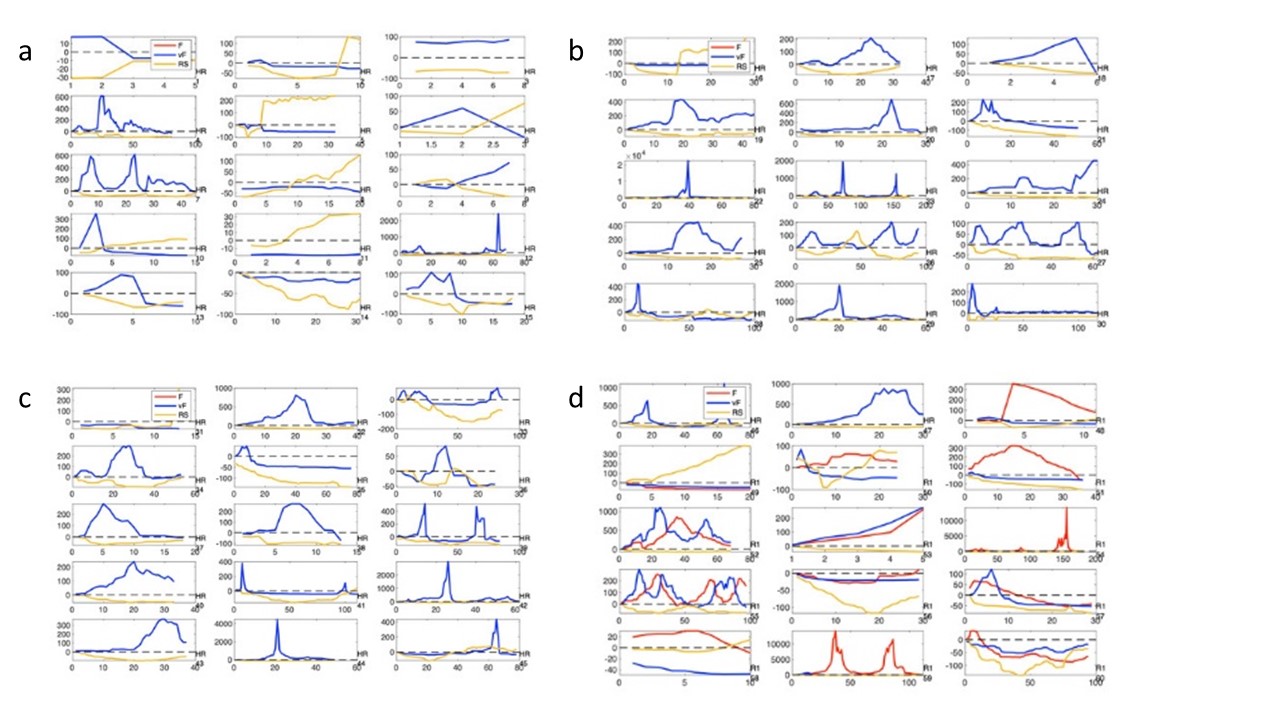

### Fig. S6e-h

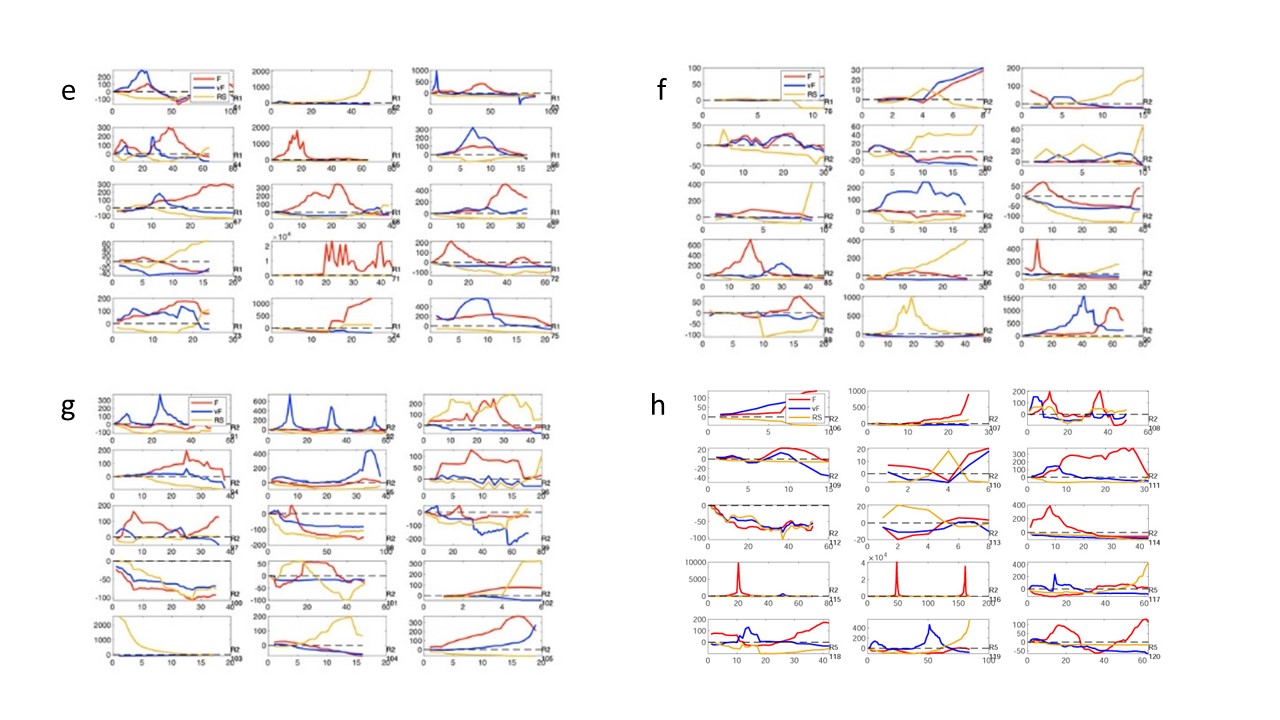
